# Unveiling transposable element expression heterogeneity in cell fate regulation at the single-cell level

**DOI:** 10.1101/2020.07.23.218800

**Authors:** Jiangping He, Isaac A. Babarinde, Li Sun, Shuyang Xu, Ruhai Chen, Yuanjie Wei, Yuhao Li, Gang Ma, Qiang Zhuang, Andrew P. Hutchins, Jiekai Chen

**Author notes:** Corresponding author. (A.P.H), (J.C.).

## Abstract

Transposable elements (TEs) make up a majority of a typical eukaryote’s genome, and contribute to cell heterogeneity and fate in unclear ways. Single cell-sequencing technologies are powerful tools to explore cells, however analysis is typically gene-centric and TE activity has not been addressed. Here, we developed a single-cell TE processing pipeline, scTE, and report the activity of TEs in single cells in a range of biological contexts. Specific TE types were expressed in subpopulations of embryonic stem cells and were dynamically regulated during pluripotency reprogramming, differentiation, and embryogenesis. Unexpectedly, TEs were expressed in somatic cells, including human disease-specific TEs that are undetectable in bulk analyses. Finally, we applied scTE to single cell ATAC-seq data, and demonstrate that scTE can discriminate cell type using chromatin accessibly of TEs alone. Overall, our results reveal the dynamic patterns of TEs in single cells and their contributions to cell fate and heterogeneity.

## Introduction

Transposable elements (TEs) are a heterogeneous collection of genomic elements that have at various stages invaded and replicated extensively in eukaryotic genomes. The vast majority of TEs are fossils, and can no longer duplicate themselves, but they remain inside the genome and in mammals occupy nearly half the total DNA^1^. Intriguingly, it is becoming clear that both the active and remnant TEs are participating in evolutionary innovation and in biological processes^2-5^, such as embryonic development^6-9^, and in human disease and cancer^10,11^. Additionally, TEs carry cis-regulatory sequences and their duplication and insertion can reshape gene regulatory networks by redistributing transcription factor (TF) binding sites and evolving new enhancer activities^12-14^. TEs transcription also has a key influence upon the transcriptional output of the mammalian genome^15^. However, the role of TEs in cell type heterogeneity and biological processes has only recently begun to be explored in depth.

Single cell RNA-seq (scRNA-seq) has developed as a powerful tool to observe cell activity^16-18^. Many new techniques have been developed to recover or reconstruct missing observations, such as spatial, temporal, and cell lineage information. However, an important source of genomic information has so far been overlooked in single cell studies: the effect of TEs. Despite their importance, we lack quantitative understanding of how those genomic elements are involved in cell fate regulation at the single cell level. As TEs pose unique challenges in quantification, due to their degeneracy and multiple genomic copies, a prerequisite to understand TEs at the single cell level is a tool to quantify the hundreds to millions of copies of repetitive elements within the genome. To this end, we developed scTE, an algorithm that quantifies TE expression in single-cell sequence data.

We firstly demonstrate scTE’s capabilities through an analysis of mouse embryonic stem cells (mESCs), which is one of the best characterized models for TE expression, as the expression of the endogenous retrovirus (ERV) MERVL marks a small population of cells in embryonic stem cell (ESC) cultures that are totipotent^19,20^, scTE could accurately recover the expected pattern of heterogeneous MERVL expression. Then, we applied our approach to several biological systems including human *in vitro* cardiac differentiation, mouse gastrulation, adult mouse somatic cells, the induced pluripotent reprogramming process and human disease data. Overall, we unveil hitherto unknown insights into complex TE expression patterns in mammalian development and human diseases.

## Results

### Quantification of TE expression in single cells with scTE

Analysis of TEs pose special challenges as they are present in many hundreds to millions of copies within the genome. A common strategy in regular analyses is to discard multiple mapped reads, however this leads to loss of information from TEs ^21^. Assigning these reads to the best alignment location is the simplest way to resolve TE-derived reads, but it is not always correct for individual copies^21,22^. To solve this problem, we designed an algorithm in which TE reads are allocated to TE metagenes based on the TE type-specific sequence. We built a framework named scTE with this strategy, scTE maps reads to genes/TEs, performs barcode demultiplexing, quality filtering, and generates a matrix of read counts for each cell and gene/TE (Fig. 1a and Supplementary Fig. 1a). scTE is easy to use, and its output is designed to be easily integrated into downstream analysis pipelines including, but not limited to, Seurat and SCANPY^23,24^. The algorithm can in principle be applied to infer TE activities from any type of single-cell sequencing based data, like single-cell ATAC-seq data, DNA methylation, and other single-cell epigenetic data.

**Fig. 1.**
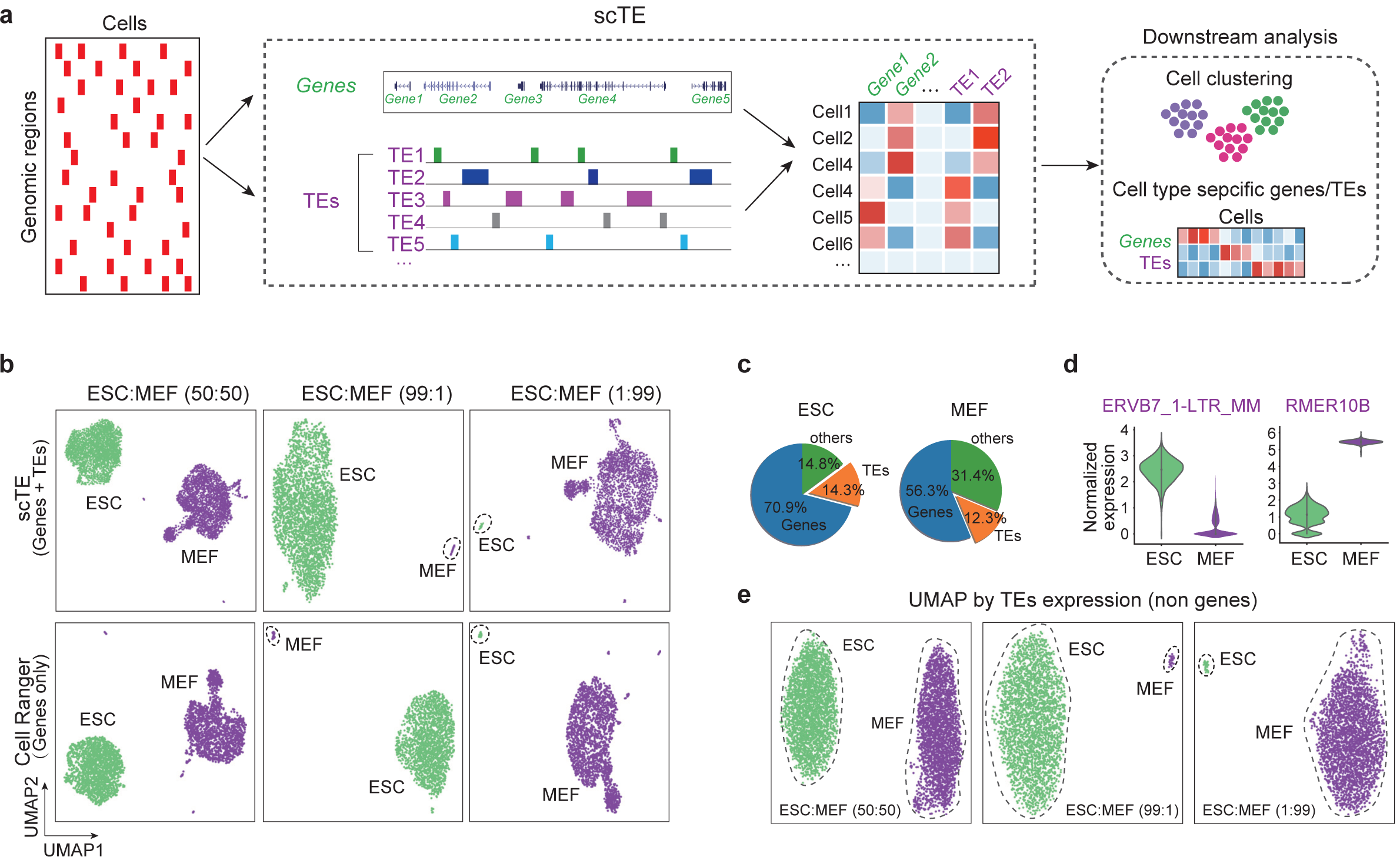
scTE workflow and applications. (**a**) Schematic of the workings of scTE. For scRNA-seq data the reads are mapped to the genome, and assigned to either a gene, or a metagene model of a TE. Multimapping read data will assign the best mapping read to a type of TE. Reads are always mapped to a gene first, and then a TE if no gene is found. The resulting assignments are then collapsed into a matrix of read counts for each cell, versus each gene/TE. This matrix can be used in downstream applications. (**b**) UMAP plot showing mixtures of MEFs and ESCs in the indicated ratios. The top panels show scTE analysis, the lower panels show Cell Ranger analysis results. Cells and are colored by their sample of origin. (**c**) Percentage of reads mapping to genes, TEs or other regions of the genome in MEFs and ESCs. (**d**) Violin plot showing the expression of selected TEs in MEFs and ESCs. (**e**) As in panel b, but only TE expression was used.

We first tested scTE’s ability by *in silico* mixing two cells lines, MEFs (mouse embryonic fibroblasts) and ESCs in different ratios^25^. Comparison with the gene-based Cell Ranger pipeline^26^, scTE shows nearly identical topology in a UMAP (Uniform Manifold Approximation and Projection) plot, and in marker genes expression (Fig. 1b and Supplementary Fig.1b). Even when one cell type only contributes a 1% minority in the mixture, scTE identified it correctly (Fig. 1b), indicating that scTE did not influence the global analysis of gene expression. These results demonstrate the sensitivity of scTE.

Next, we sought to explore TE expression, around 12-14% of the reads were derived from TEs (Fig. 1c). Requiring at least 2-fold change and FDR<0.05, scTE detected 150 significantly differentially expressed TEs between ESCs and MEFs (Supplementary Fig. 1c), including ERVB7_1-LTR_MM, which is highly expressed in ESCs, and RMER10B in MEFs (Fig. 1d and Supplementary Fig.1d). Furthermore, UMAP based on single cell TE expression alone could distinguish the cell types with the expected ratio (Fig. 1e), demonstrating TE expression discerns cell identity.

### Deciphering TE heterogeneity in mouse ESCs and during human cardiac differentiation

It is known that a small subset of ESCs acquire a totipotent state named 2C-like cells and express a MERVL TE which also marks the embryonic 2-cell stage^19,27,28^. scTE could correctly identify this rare 2C-like subpopulation in UMAP plots, based on the specific marker genes *Zscan4c* and *Tcstv3*, and the expression of MERVL and MT2_Mm TEs (Fig. 2a, b and Supplementary Fig. 2a, b)^19,29^. If we discarded multiple mapped reads and only considered unique reads, the level of MERVLs was reduced, but it was still specifically expressed in the 2C-like cells (Supplementary Fig. 2c). This confirms that scTE can correctly identify known TE patterns.

**Fig. 2.**
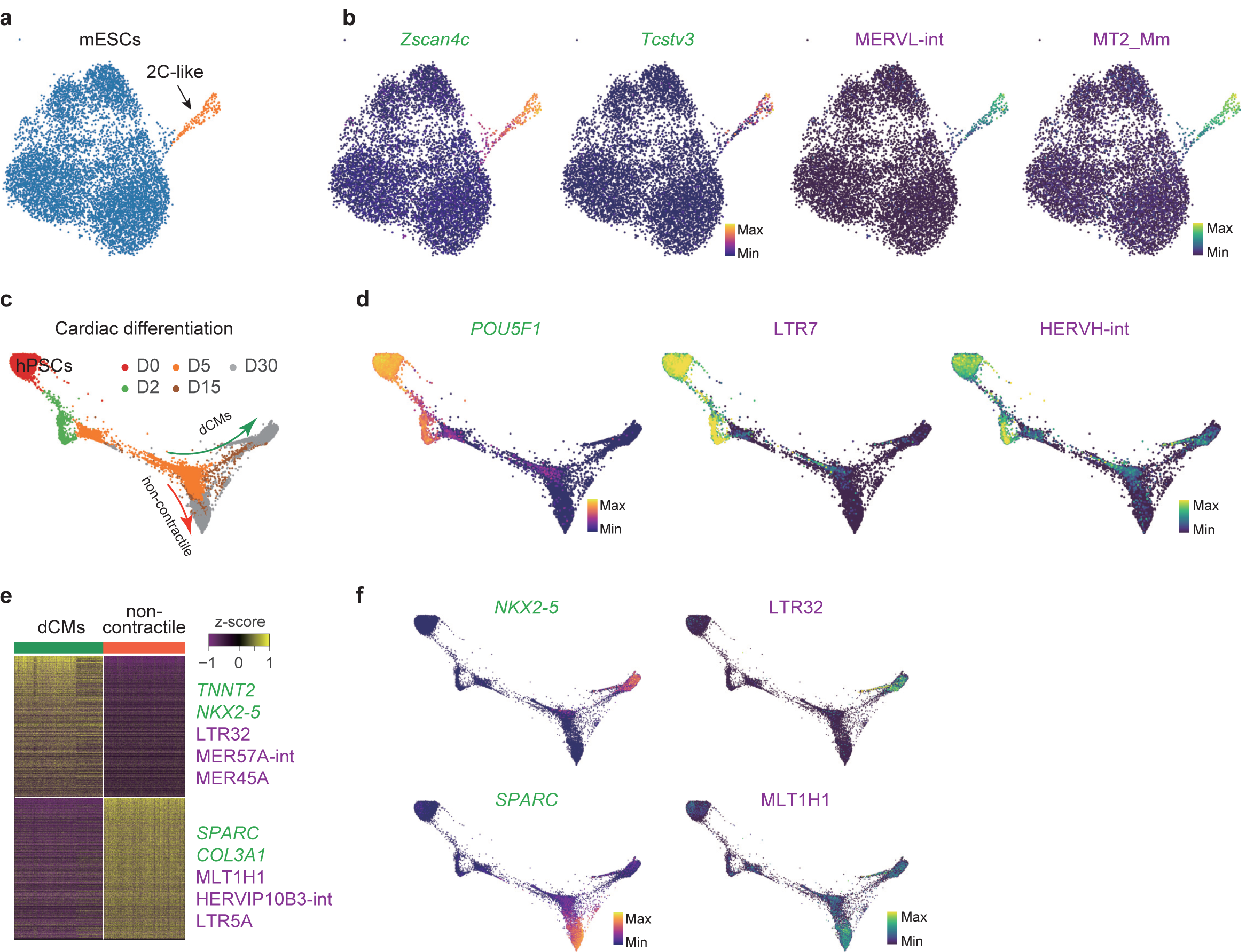
Dynamic transcription of TEs in ESCs and during cardiac differentiation. (**a**) UMAP plot of mouse ESCs. Cells are colored by cell type cluster. (**b**) Same as panel a, but cells are colored based on the expression of the indicated genes and TEs. *Zscan4c* and *Tcstv3* are marker genes for the 2C-like cells. (**c**) Trajectory reconstruction of single cells through a cardiac differentiation timecourse showing the definitive cardiomyocytes (dCMs) branch and non-contractile branch. Days of differentiation (D) are labelled. (**d**) As in panel c, but cells are colored by the expression of the indicated genes and TEs. (**e**) Heatmap of expression differences between dCM (contractile) branch and non-contractile branch cells, selected differentially expressed genes and TEs are labelled. (**f**) As in panel d, but cells are colored by the expression level of the indicated genes and TEs.

In humans, HERV-H LTRs are expressed in early embryos and human pluripotent stem cells (hPSCs), and contribute to pluripotency maintenance and somatic reprogramming^6,30-32^, but little is known about TE expression dynamics during differentiation to somatic cells. Applying scTE to an scRNA-seq time series of hPSCs differentiating to cardiomyocytes^33^, we accurately recovered the repression of HERV-H LTRs including LTR7 and HERVH-int during differentiation, concomitant with reduction in the expression of the pluripotency factor *POU5F1* (Fig. 2c, d and Supplementary Fig. 2d). During *in vitro* cardiac differentiation of hPSCs there is a bifurcation towards definitive cardiomyocytes (dCM) and non-contractile cells (Fig. 2c). Between these two branches, marked by *NKX2-5* and *SPARC*, respectively, we found differential expression of TEs such as LTR32, MER57A-int and MER45A in the dCM cells, whilst, MLT1H1, HERVIP10B-int and LTR5A were specifically expressed in the non-contractile cells (Fig. 2e, f and Supplementary Fig. 2e). Independent bulk RNA-seq data^34^ demonstrated that these TEs were expressed in late cardiac differentiation (Supplementary Fig. 2f), however, as the bulk is a mixture of dCM and non-contractile cells, the restriction of these TEs to divergent fates can only be observed in the scRNA-seq data. This highlights the importance of analyzing TE expression in sc-RNA-seq data, as MLT1H1 is very high in the bulk RNA-seq, but this hides the reality that it is restricted to the non-contractile cells and plays no role in dCMs (Fig. 2e, f and Supplementary Fig. 2f).

### Analysis of TEs in mouse gastrulation and early organogenesis reveals the widespread cell fate-specific expression of TEs

The previous analysis showed how TE expression contributed to *in vitro* cardiac differentiation, next we explored complex *in vivo* developmental processes. TE expression is dynamic during pre-implantation development^6^, however the expression of TEs in gastrulation has not been described. We took advantage of the single-cell time course of mouse gastrulation^16^. Analysis with scTE did not introduce any unexpected sample-bias, and a side-by-side comparison could retrieve similar patterns of marker gene expression in the expected lineages (Fig. 3a and Supplementary Fig. 3a-f). We found every lineage expressed a series of lineage-specific TEs (Fig. 3a, b, and Supplementary Fig. 4a-c). In the extraembryonic ectoderm cells, IAP and RLTR45-family TEs were activated (Fig. 3b, c), and in *Apoa2*+ extraembryonic endoderm cells, MER46C, RLTR20B3 and LTRIS2 were up-regulated (Fig. 3b, d). The expression of these TEs was validated using bulk RNA-seq from *in vitro*^35-37^ mimics of these embryonic stages including ESCs, epiblast stem cells (EpiSCs), extraembryonic endoderm cells (XENs) and trophoblast stem cells (TSCs) (Fig. 3e). Other embryonic lineages, particularly the *Gypa*+ erythroid and the *Tnnt2*+ cardiomyocyte lineages expressed specific TEs such as L1_Mur and L1ME3D, respectively (Fig. 3b, f).

**Fig. 3.**
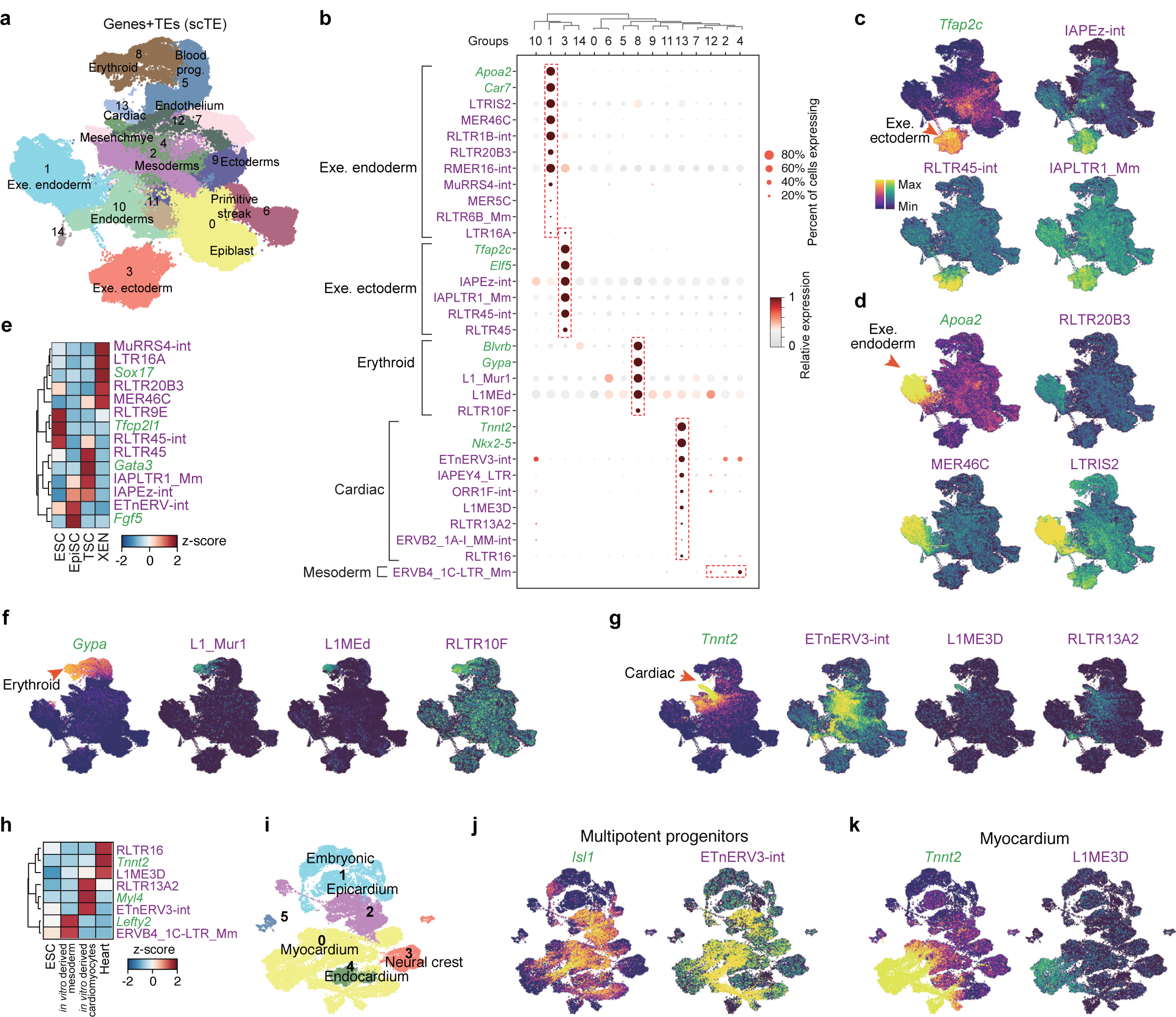
Widespread cell type-specific expression of TEs during gastrulation. (**a**) UMAP plots of the mouse gastrulation data using both genes and TEs. Selected lineages are labelled (Leiden, resolution=0.3). (**b**) Dot plot showing a selection of marker genes and TEs for the indicated cell lineages. (**c**) Expression of the indicated extra embryonic ectoderm gene *Tfap2c* and selected TEs. (**d**) Expression of the extra embryonic endoderm marker gene *Apoa2* and selected TEs. (**e**) Expression of the indicated TEs and marker genes in bulk RNA-seq data from ESCs, EpiSCs, XEN (extra embryonic endoderm cells) and TSCs (trophoblast stem cells). *Tfcp2l1, Fgf5, Gata3* and *Sox17* serve as markers for ESCs, EpiSCs, TSCs, and XEN cells, respectively. Data is displayed as a z-score using the variance from all genes. (**f**) Expression of the erythroid marker gene *Gypa*, and selected TEs. (**g**) Expression of the cardiac marker gene *Tnnt2* and selected TEs. (**h**) Expression of the indicated TEs and marker genes from bulk RNA-seq data. (**i**) UMAP plot of the embryonic mouse heart scRNA-seq data using both TEs and genes. The indicated developmental stages are labelled as in the original study. (**j-k**) UMAP as panel i, but cells are colored by the expression of indicated genes/TEs.

As this dataset provides dynamic trajectories for each lineage, we wondered if TEs where transiently activated during cell fate commitment. To this end, we noticed ETnERV3-int, whose expression coincides with the early development of the cardiac fate from the mesoderm, and is reduced in *Tnnt2*+ cells, while L1ME3D was expressed in the *Tnnt2*+ cells (Fig. 3g). Consistently, ETnERV3-int was specifically expressed in *in vitro* derived cardiomyocytes, which more closely resemble a fetal state, whilst L1ME3D was expressed only in the mature heart (Fig. 3h)^38,39^. However, the bulk samples could not capture the complexity of the transient expression of ETnERV3-int which extended from the late epiblast into the endoderm and mesoderm. To expand on this, we reanalyzed an scRNA-seq dataset of the developing mouse embryonic heart^40^ (Fig. 3i and Supplementary Fig. 5a-c), and found that ETnERV3-int was expressed in the myocardium and epicardium, but not in the endocardium, neural crest and embryonic cells (Fig. 3j). L1ME3D was expressed in *Tnnt2*+ myocardium, however in an inverse pattern with respect to ETnERV3-int (Fig. 3j, k). Therefore, ETnERV3-int activity is present in an intermediate stage in cardiac lineage development. Intriguingly, there was a close relationship between the expression of ETnERV3-int and *Isl1* gene, which marks multipotent progenitors ^40^ (Fig. 3j). These results highlight the complex patterns of TE expression in developmental processes.

### Widespread tissue-specific expression of TEs in somatic cells

TE activity is considered to be silenced in somatic cells except LINE-1 expression and retrotransposition in the developing brain^27,28,41^. As we revealed unexpected heterogeneity of TEs in somatic MEFs and during organogenesis, we next measured TE expression in somatic cells using the Tabula Muris large scale scRNA-seq dataset that profiles 20 mouse organs^42^ (Fig. 4a). Surprisingly, our analysis revealed in total 130 TEs that were specifically expressed in distinct cell types (Fig. 4b and Supplementary Fig. 6a). These associations include the expected expression of LINE1 elements in brain cells, of which many L1 family members like L1MEh, L1M, L1MC4a, L1MA7 and L1P5 elements are specifically expressed in oligodendrocytes or microglia (Fig.4c and Supplementary Fig. 6a). We also found expression of LTR58, MLT1EA-int, MER110 and RLTR46 that specifically in B cells, T cells, type B pancreatic cells and hepatocytes, respectively (Fig.4c). Next, we took advantage of the Tabula Muris dataset to measure overall TE expression heterogeneity, and, in general, the LTRs and DNA transposons are the major source of heterogeneity (Fig. 4d, e).

**Fig. 4.**
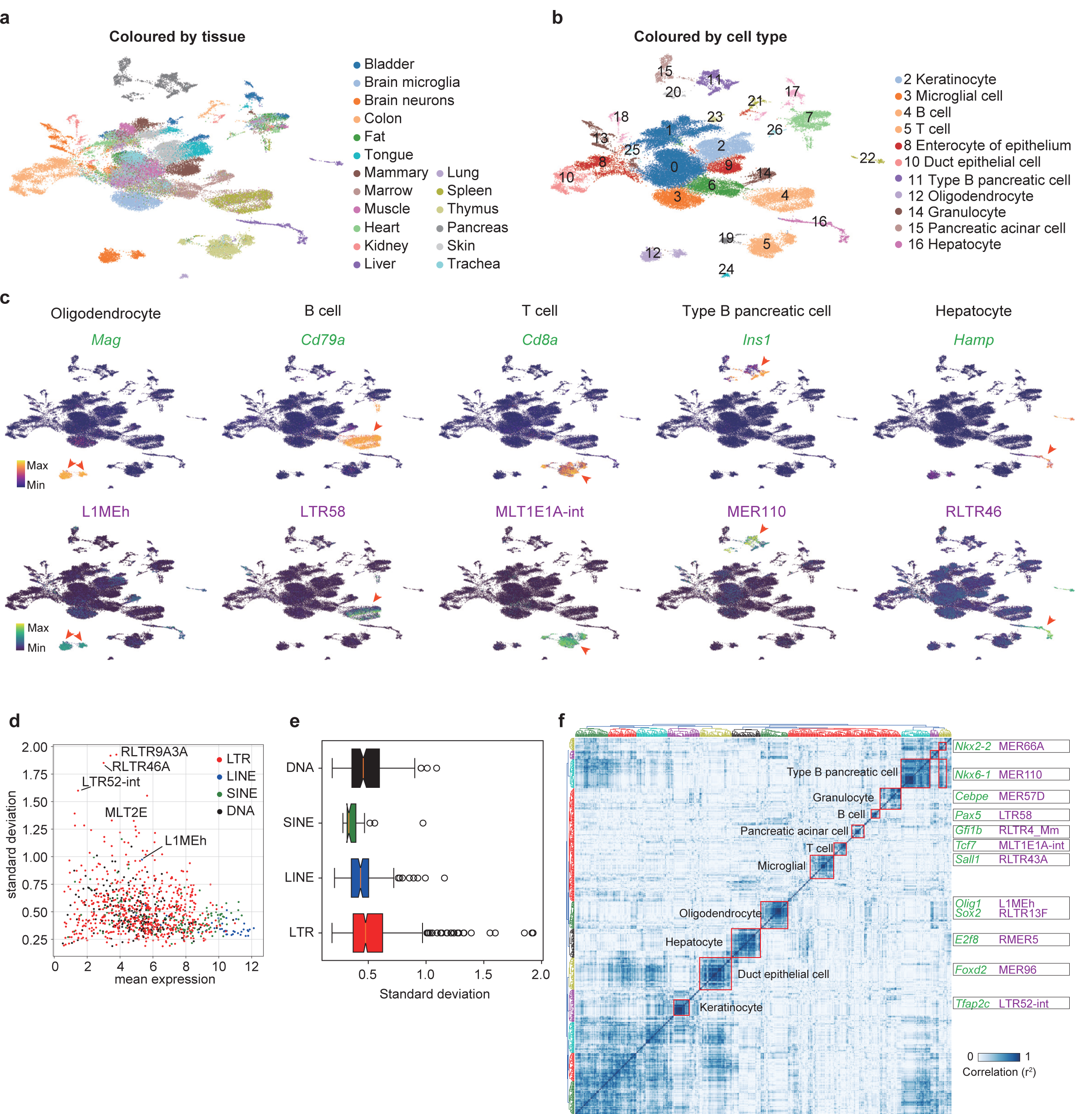
Class-specific expression of TEs in somatic cells. (**a**) UMAP plots of the Tabula Muris data, using both genes and TEs as analyzed with scTE. The tissue sources for the cells is indicated. (**b**) UMAP plot as in panel a, but clustered into groups (Leiden, resolution=0.5). (**c**) Same as panel b, but cells are colored by the expression of indicated genes/TEs. (**d**) Scatter plot showing TE expression heterogeneity. The x-axis is the mean expression for cells from panel b, the y-axis is the standard deviation for each TE type, the higher standard deviation represents higher heterogeneity across cell types. (**e**) Boxplot for the standard deviations for each class of TEs. (**f**) Correlation heatmap showing the co-expression of TFs and TEs.

TE expression is regulated by chromatin modification and transcription factors (TFs) ^3^, thus, we wondered if we could infer the regulatory network between TFs and TEs from large scale scRNA-seq data, taking advantage of the improved cell type definitions from the scRNA-seq data. The co-expression relationships often reflect biological processes in which many genes with related functions are coordinately regulated. Therefore, we reasoned that if a TE is regulated by a TF, they should be co-expressed. To identify TF-TE regulatory relationships, we performed co-expression analysis, and revealed the specific co-clustering of neural genes and TEs (*Sox2* and *Olig1*), the immune system (*Cebpe, Tcf7, Pax5* and *Sall1*), the endoderm/pancreas (*Gfi1b, Nkx6-1* and *E2f8*), and other lineages (Fig. 4f and Supplementary Fig. 6b). Motif analysis also showed that the SOX2 motif was significantly enriched within RLTR13F TEs (Supplementary Fig. 6c). These results highlight the deep link between TE and TF activity Indicating those TFs may be responsible for activating TEs in the corresponding cell types.

We next explored in closer detail neural and immune cell lineages as TE activity is known to regulate neural activity and immune responses ^43-45^. Subgrouping the cells from microglia and neuron samples identified several distinct cell types (Supplementary Fig. 7a-c), within which cell type-specific expression of TEs was observed (Supplementary Fig. 7d, e). Next, with the pooled immune cells from marrow, spleen and thymus, 12 distinct immune cell subtypes were defined (Supplementary Fig. 7f, g). Intriguingly, besides finding additional cell type-specific TEs in T cells, B cells and granulocytes, a series of TEs were restricted to subtypes of T cells and B cells (Supplementary Fig. 7h, i,). These data show different degrees of subtype specific signatures of TEs in the neural and immune system, and highlight the importance of looking beyond only genes when exploring how those systems differ.

### TEs are activated during somatic cell reprogramming, in a heterogonous and cell branch restricted manner

The above analysis has revealed the well-ordered dynamic expression of TEs in developmental processes, we then wondered if TEs undergo similar stage-specific regulation during somatic reprogramming. Somatic cells can be reprogrammed to induced pluripotent stem cells (iPSCs) by various methods, such as ectopic expression of a group of pluripotency transcription factors^25,46,47^, or cocktails of chemicals^48,49^. The reprogramming process is highly heterogeneous, with abundant non-reprogramming cells and divergent cell fate transition routes^25,50^. We took advantage of reprogramming scRNA-seq data to investigate the activity of TEs during these drastic cell fate transitions. Reprogramming induced by *Oct4/Pou5f1, Klf4, Sox2* and c-*Myc* (OKSM) generates detectable intermediate branches, including iPSCs, trophoblast, stromal and neural-like cells (Fig. 5a and Supplementary Fig. 8a-d)^50^. We identified specifically expressed TEs in each cell branch (Supplementary Fig. 8a-d). For example, the TEs ERVB7_1-LTR_MM, IAPEz-int, RLTR4_Mm, and Lx were specifically expressed in iPSCs, trophoblast, stromal and neural-like branches, respectively (Fig. 5b). ERVB7_1-LTR_MM (MusD) and IAPs are up-regulated during reprogramming ^51^, however using scRNA-seq data we show that only ERVB7_1-LTR_MM, as well as ETnERV-int and RLTR13G, were up-regulated in the successful reprogramming route, initiating at the mesenchymal-to-epithelial transition (MET) and peaking at the iPSCs stage (Fig. 5b and Supplementary Fig. 8a). In contrast, the trophoblast-branch expressed IAPEz-int and IAPLTR1_Mm (Fig. 5b and Supplementary Fig. 8c), which are also expressed in *in vivo* extra embryonic ectoderm cells (Fig. 3c), suggesting consistent regulation between development and reprogramming.

**Fig. 5.**
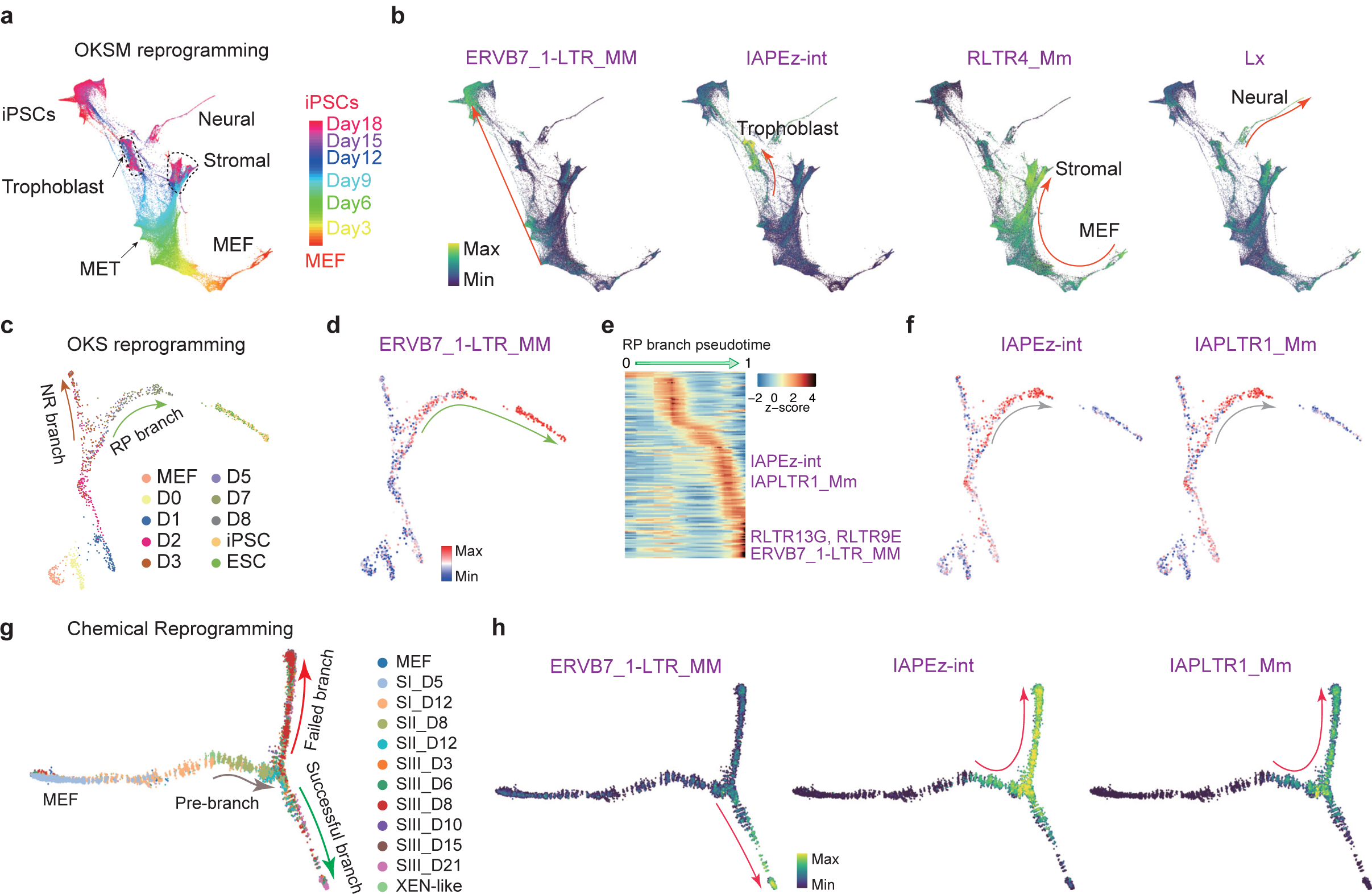
Stage-specific expression of TEs in somatic cell reprogramming. (**a**) Trajectory reconstruction during OKSM reprogramming, cells are colored by time point. (**b**) As in panel a, but cells are colored by the expression of the indicated TEs. (**c**) Force-directed (FR) layout of cells during OKS reprogramming, cells are colored by time point. (**d**) Same with panel c, but cells are colored by the expression change of the ERVB7_1-LTR_MM TE during reprogramming. (**e**) Expression heatmap of the top 145 dynamically expressed TEs in a pseudotime ordering for the RP branch, selected TEs are indicated. (**f**) Expression changes of the indicated TEs during reprogramming. (**g**) Trajectory reconstruction during chemical reprogramming, cells are colored by time point. (**h**) As in panel g, but showing TE expression specific to the successful or failed branches of reprogramming.

We then analyzed reprogramming induced by *Oct4, Klf4*, and *Sox2* (OKS) ^25^ or only chemicals^29^. There are two validated branches during OKS-mediating reprogramming^25^ (Fig. 5c), and we found many TEs, such as ERVB7_1-LTR_MM, that were specifically up-regulated in the reprogramming-potential (RP) branch, and were excluded from the non-reprogramming branch (Fig. 5d and Supplementary Fig. 8e). IAPEz-int and IAPLTR1_Mm were expressed in the RP branch but were ultimately silenced in the reprogrammed cells (Fig. 5e, f), suggesting IAPs were only activated in a pre-reprogrammed state and may impede the final step of pluripotency acquisition. We validated the expression of ERVB7_1-LTR_MM and IAPs by qRT-PCR (Supplementary Fig. 8f), demonstrating that IAPs are silenced in ESCs. Similar to OKS-mediated reprogramming, chemical-mediated reprograming bifurcates into two branches (Fig. 5g and Supplementary Fig. 8g)^29^, and TEs, marking an intermediate 2C-like program, were activated at the root of the successful branch (Supplementary Fig. 8h, i). ERVB7_1-LTR_MM and RLTR13G were specifically up-regulated in the successful branch, whilst IAPEz-int and IAPLTR1_Mm were activated in the pre-branch and failed branch (Fig. 5h and Supplementary Fig. 8j, k).

The similar expression pattern of TEs among the three distinct reprogramming systems described above, suggests there are common regulatory mechanisms. Indeed, we found IAPLTR1_Mm TEs are rich in DNA-binding motifs for JUN and IRF2 (Supplementary Fig. 8l), whose expression closely matched IAP expression in all three reprogramming systems (Supplementary Fig. 8m) and are known to impair reprogramming^52,53^. This suggests that their downregulation deactivates the IAPs before the finalization of reprogramming, indicating IAPs may impede the final step of reprogramming. Overall, these results indicate TEs have a deeper unappreciated role in iPSC formation.

### Inferring TE Associated Accessibility from scATAC-seq Data

Beyond scRNA-seq, many other single-cell sequencing techniques^54-56^ have shown great potential to explore cell heterogeneity and increased insight could be fueled by the additional information provided by scTE. For instance, we reasoned that scTE would be informative for the analysis of scATAC-seq data and potentially other single-cell epigenetic data because TEs have a wide array of chromatin states^3^, are widely bound by transcription factors^57^, and can act as enhancers^14^ (Fig. 6a). We then applied scTE to a dataset of fluorescence-activated cell sorted (FACS) mouse cells^58^, including cardiac progenitor cells (CPCs), CD4^+^ T cells, ESCs and skin fibroblasts (SFs). Intriguingly, scTE could accurately recover the expected cell types, based on only the reads that mapped to TEs (Fig. 6b). Specific accessibility of RLTR13A, RLTR4_Mm, RLTR13G and RMER19B/C was found in the CPCs, CD4+ T cells, ESCs and SFs, respectively (Fig. 6c, d and Supplementary Fig.9a). And motif enrichment of these cell-type specific TEs revealed known master regulators of these cell types, such as GATA4/HAND1/T for CPCs, ETS1/TCF3 for T cells, SOX2/POU5F1/NR5A2 for ESCs and FOS/MAF for SFs (Supplementary Fig. 9b), indicating these TEs may act as cis-regulatory elements bound by transcription factors. For instance, scTE reveals there is an RLTR13A TE within an intron of *Smyd1*, a gene essential for heart development^59-61^, which was specifically open in CPCs (Fig. 6e), and was specifically expressed in the myocardium of the fetal heart (Fig. 6f). Applying scTE to scATAC-seq data of peripheral blood monocyte cells (PBMC) was also able to recover the major cell types and cell type-specific TEs (Supplementary Fig. 8c-f), which can be validated by independent bulk ATAC-seq data from FACS sorted cells (Supplementary Fig. 8g)^62^. These results indicate that quantifying chromatin accessibility on TE regions is informative for characterizing cell types and may assist the problems posed by scATAC-seq analysis due to its especially sparse nature^63^.

**Fig. 6.**
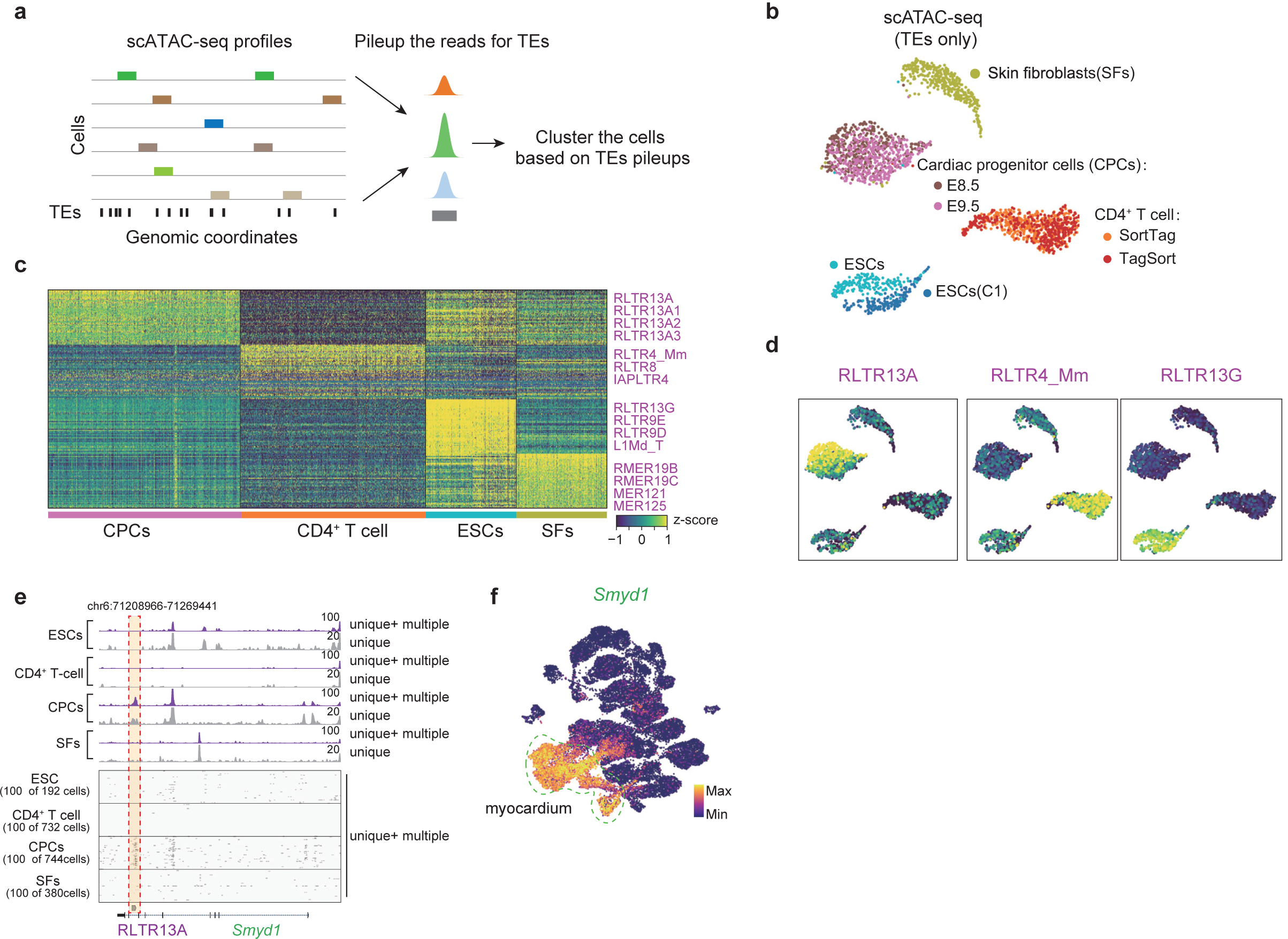
Analysis of the Chromatin State of TEs in Single-Cell ATAC-seq data. (**a**) Schematic plot of scTE for scATAC-seq data analysis. The reads are mapped to the genome, and assigned to a metagene TE, and then the cells were clustered based on the TE matrix. (**b**) UMAP plot of the TE chromatin state from scATAC-seq data for a selection of FACS-purified mouse cell types. (**c**) Heatmap of the top 50 cell type-specific opened TEs in the indicated cell types, selected example TEs are indicated. (**d**) UMAP plot as in panel b, but cells are colored by chromatin-state of the indicated TEs. (**e**) Genome tracks showing the aggregate scATAC-seq profiles (top panel). Randomly selected 100 single cell profiles are show below the aggregated profiles (bottom panel). With include (unique + multiple) or exclude (unique) multiple mapped reads. (**f**) UMAP plot of the expression of the myocardium marker gene *Smyd1*, from the cardiogenesis data, see Fig. 3i.

### Disease-specific expression of TEs

The unexpected widespread TE heterogeneity amongst embryonic and somatic cell types and cell fate transitions raised the question as to whether there is TE heterogeneity in diseased cells. Alzheimer’s disease (AD) is an age-associated neurodegenerative disorder that is characterized by progressive memory loss and cognitive dysfunction for which there is no known cure. TEs have been reported to be highly active during aging and may contribute to age-dependent loss of neuronal function^64^. To explore the expression of TEs in AD, we reanalyzed the scRNA-seq data from a mouse model of AD expressing five human familial AD gene mutations, which contained 13,114 single cells with age and sex-matched wild-type (WT) controls using the MARS-seq platform^65^ (Fig. 7a). Projecting the cells with a UMAP, we recovered the major groups of cells in AD and WT, including the unique disease-associated microglia cluster cells (M2) identified in the original study (Fig. 7b and Supplementary Fig. 10a). Differential expression analysis demonstrated significant changes in gene expression in M2, including previously described AD risk factors such as *Apoe, Tyrobp, Lpl, Cstd* and *Trem2* (Fig. 7c and Supplementary Fig. 10b). Intriguingly, we also found many TEs such as ERVB7_2-LTR_MM, RLTR17, RLTR28 and Lx4B that were significantly higher and specifically expressed in M2 (Fig. 7c, d and Supplementary Fig. 10c), indicating those TEs may also be involved in AD development.

**Fig. 7.**
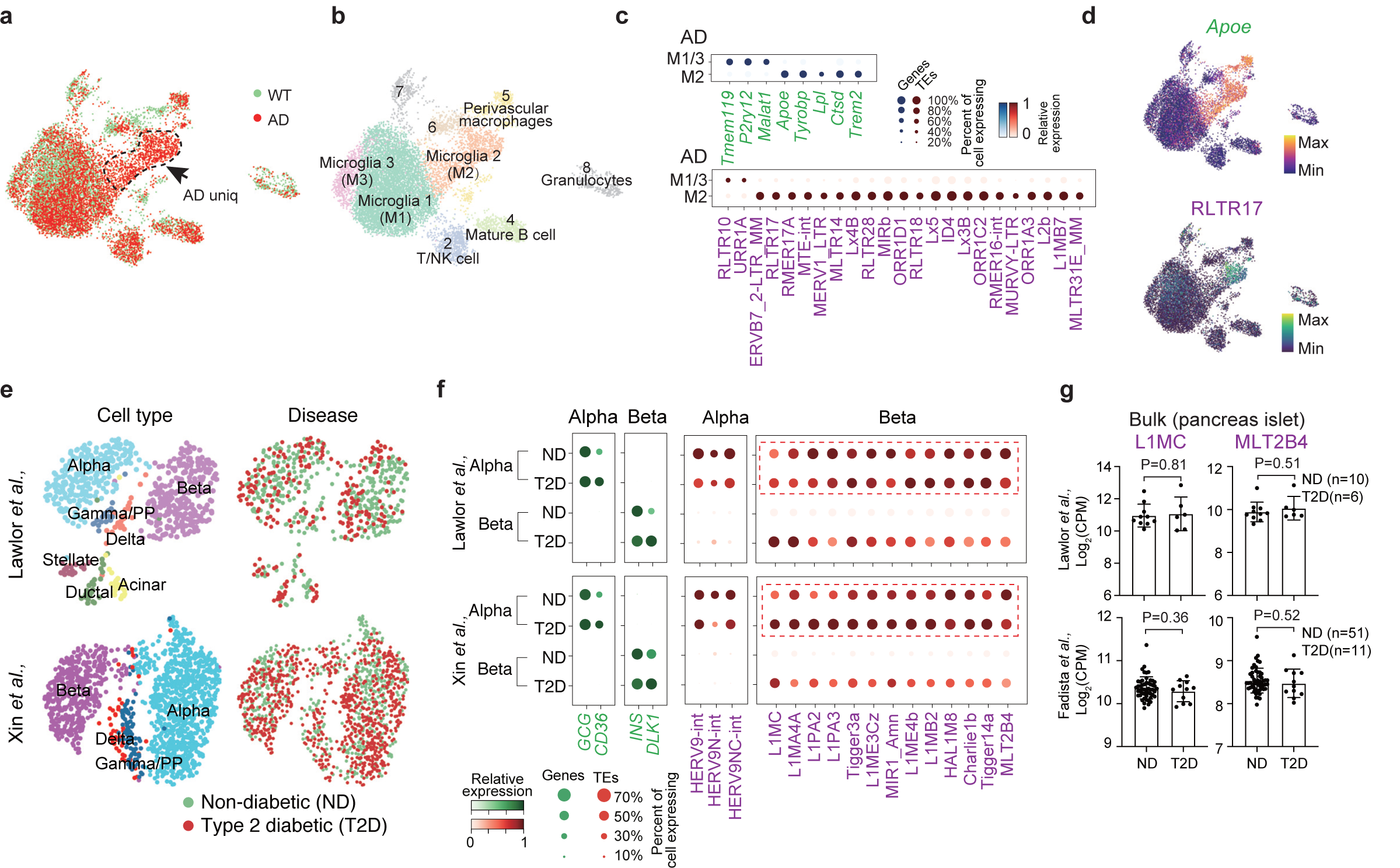
TEs are differentially expressed in single cells in the diseased state. (**a**) UMAP plot of the single cells genes and TE expression, cells are colored by WT (wild-type) and AD (Alzheimer’s disease) state. (**b**) UMAP plot, as in panel a, but clustered into groups (Leiden, resolution=0.5). (**c**) Dot plot showing the differential expressed genes (top) and TEs (bottom) between disease associated microglia (M2) and homeostatic microglia (M1/3) in AD mice. (**d**) UMAP plot, as in panel a, but cells are colored by the expression of the indicated *Apoa2* or the TE RLTR17. (**e**) UMAP plots of pancreatic islet cells. Cells are colored by cell types (left) or disease-state (right). Cell types were annotated according to the metadata from the original study, and matched the expression of known marker genes. (**f**) Dot plot showing marker gene expression (green) or TEs (red) differentially expressed between healthy and T2D alpha and beta cells (Benjamini-Hochberg corrected Wilcoxon rank-sum test, P<0.01, and at least >2-fold change between groups). (**g**) Bar charts showing the expression of the indicated TEs from bulk RNA-seq data. P-value was from an unpaired t-test.

Type 2 diabetes (T2D) is a common human disease caused by a combination of increased insulin resistance and reduced mass or dysfunction of pancreatic beta cells. We reanalyzed scRNA-seq from two independent studies of the human pancreas in healthy and T2D individuals^66,67^. The major cell types in the pancreas, including alpha, beta, gamma/PP and delta cells clustered without a visible disease-specific pattern, indicating no drastic change in cell type (Fig. 7e and Supplementary Fig. 10d). Contrasting the transcriptome from healthy and T2D in each cell type independently, *CD36* and *DLK1* was up-regulated in T2D alpha and beta cells respectively (Fig. 7f), as reported by the original studies^66,67^. Notably, many TEs were significantly highly expressed in T2D beta cells, including L1MC, L1MA4A, Tigger3a, MLT2B4. This differential expression pattern was near identical between the two independent datasets (Fig. 7f). Critically, none of these observations could be observed using bulk RNA-seq datasets (Fig. 7g and Supplementary Fig. 10e)^66,68^, which might be due to the high expression of these TEs in both normal and T2D alpha cells, emphasizing the importance of analysis at single-cell resolution.

As a final human disease dataset we reanalyzed a glioblastoma scRNA-seq experiment^69^, and were able to identify TEs specifically expressed in neoplastic cells and that were correlated with the expression of *EGFR* (Supplementary Fig. 10f-h), a gene upregulated in a large percentage of glioblastomas^69^. Above all, these results revealed the dysregulation of TE expression in diseased human cells, which deserves further mechanistic study and may help to identify new diagnostic markers and therapeutic targets.

## Discussion

TEs are the most abundant elements in the genome, however, the understanding of their impact on genome evolution, function and disease remains limited. The rise of genomics and large-scale high-throughput sequencing has shed light on the multi-faceted role of TEs. However, many genomic studies exclude TEs due to difficulties in their analysis as a consequence of their repetitive nature^21^. Thus, TE analysis often requires the use of specialized tools to extract meaning^5,22^. Here, we developed scTE specifically for the analysis of TEs from single-cell sequencing data. By taking advantage of this tool we could recover previously identified phenomena such as MERVL and LTR7/HERVH expression in mouse and human ESCs, respectively. We then revealed widespread heterogeneity of TE expression throughout embryonic development, in mature somatic cells, during the reprogramming process and in human diseases, and discovered a wealth of cell fate-specific TE expression. These associations with cell fate cannot be observed when only considering bulk samples, demonstrating the enormous power of single-cell sequencing, and the importance of analyzing TE expression.

One of the key findings of our analysis has revealed the various TEs that are specifically expressed in different cell types. The expression of TEs during the pre-implantation development stage has been demonstrated previously^6^, our findings extend this to gastrulation and early organogenesis. We find a wide array of expression of TEs in the extraembryonic tissues, which may be related to their activity as enhancers^70^. Furthermore, we show the expression of TEs within the specific lineages in the developing fetal heart. In addition, TEs are also heterogeneously expressed between cell types in adult somatic cells, which has not been demonstrated before, as TEs are thought to be primarily silent in adult tissues. Notably, we found a vast of trove of TEs that are expressed in the brain and the immune system, and individual TE types that are specifically expressed in different sub cell types. Considering the close relationship between the evolution of immune system, brain and TEs^43-45^, these results hint at further functions for TEs in these two systems.

How cells decide their fate is a fundamental question in biology. Stem cell differentiation and somatic cell reprogramming are both powerful *in vitro* models that mimic *in vivo* development and have provided great insight into cell fate decisions. However, how TEs are involved in these processes is still largely unknown. In this study, we have identified the TEs LTR32 and MLT1H1 that were differentially regulated between contractile and non-contractile cell fate decisions during human cardiac differentiation. In addition, we also found ERVB7_1-LTR_Mm and IAP elements divergent expression during reprogramming, whereas ERVB7_1-LTR_Mm may promote iPSC formation, IAP elements need to be silenced at the final stage before iPSCs formation (Fig. 5b, f, h). These mechanisms are shared among the Yamanaka factor based and chemical based reprogramming systems, indicating a tight association between TEs and cell fate decisions.

Considering the growing implication that TEs are important contributors to human disease, their study is becoming increasingly important. In addition to the ability of TEs to impact genomic stability as they duplicate^71^, which has clear implications for the development of cancer^72^, TEs are also playing more subtle roles in epigenetic control and transcript expression. For example, TEs are spliced into chimeric transcripts that drive the expression of oncogenes^11^. Similarly, the expression of TEs has been associated with several nervous system-related disorders, including neurodegeneration^10^, and L1 LINE expression is important in inflammation during aging^73^. In our work, we demonstrate that in single cells of the pancreas there is substantial TE expression deregulation in the beta cells, which is suggestive of epigenetic dysfunction and a loss of control over TE expression. Critically, this observation cannot be observed from bulk pancreatic islet samples. Considering the growing importance of exploring human disease using primary patient samples, the analysis of TEs should be included. However, to date the contribution of TE expression to the aging and diseased states remains relatively unexplored. Our approach will be an important tool in understanding the contributions of TEs to cellular heterogeneity in a variety of systems and in human disease.

## Methods

### Software availability

scTE is available at https://github.com/jphe/scTE. The code is freely available and is released under the MIT license. scTE requires Python >3.6, and the python module numpy, scTE supports the Linux and Mac platforms. Software code for the analysis of the data in this manuscript can be found at: https://github.com/jphe/scTE/tree/master/example.

### scTE pipeline

The input data for scTE consists of the annotation files for genes and TEs, and alignment files in either the SAM or BAM format^74^. By default, scTE uses GENCODE^75^ and the UCSC genome browser Repeatmasker track^76^ annotations for genes and TEs, respectively. The SAM/BAM file contains the aligned read genome locations. Many alignment programs can distinguish reads that have a unique alignment in the genome (termed unique-reads) or map to multiple genomic loci (termed multimapping reads or non-unique reads). Multimapping reads are critical for TE quantification, as TEs contain many repeated sequences and non-unique reads often map inside the TEs. To get an accurate quantitation of the number of reads mapping to TEs these reads should be preserved. However, in many analysis pipelines these reads are discarded. scTE recommends aligners to keep all of the mapped reads, and we recommend that the best single aligned multimapped read be kept. The reads can be aligned by any genome aligner, but the aligned reads must be against the genome (i.e. not against a set of genes or transcript assembly). scTE is most tuned to STAR-solo^77^ or the Cell Ranger pipeline outputs, and can accept BAM files produced by either of these two programs. For other aligners, the barcode should be stored in the ‘CR:Z’ tag, and the UMI in the ‘UR:Z’ tag in the BAM file. If the UMI is missing or not used in the scRNA-seq technology (for example on the Fluidigm C1 platform), it can be disabled with – UMI False (the default is True) switch in scTE. If the barcode is missing it can be disabled with the –CB False (the default is True), and instead the cell barcodes will be taken from the names of the BAM files (multiple BAM files can be provided to scTE with the –i option).

### scTE gene and TE indices

scTE builds genome indices for the fast alignment of reads to genes and TEs. These indices can be automatically generated using the commands:

~~~
scTE_build -g mm10 # mouse genome
scTE_build -g hg38 # human genome
~~~

These two scripts will automatically download the genome annotations, for mouse:

ftp://ftp.ebi.ac.uk/pub/databases/gencode/Gencode_mouse/release_M21/gencode.vM21.annotation.gtf.gz

http://hgdownload.soe.ucsc.edu/goldenPath/mm10/database/rmsk.txt.gz

Or for human:

ftp://ftp.ebi.ac.uk/pub/databases/gencode/Gencode_human/release_30/gencode.v30.annotation.gtf.gz

http://hgdownload.soe.ucsc.edu/goldenPath/hg38/database/rmsk.txt.gz

These annotations are then processed and converted into genome indices. The scTE algorithm will allocate reads first to gene exons, and then to TEs, by default. Hence TEs inside exon/UTR regions of genes annotated in GENCODE will only contribute to the gene, and not to the TE score. This feature can be changed by setting ‘–mode/-m exclusive’ in scTE, which will instruct scTE to assign the reads to both TEs and genes if a read comes from a TE inside exon/UTR regions of genes.

### Analysis of 10x-style data

scRNA-seq data was processed using the scTE 10x pipeline, Briefly, reads were aligned to the genome using STARsolo^77^ with the setting ‘--outSAMattributes NH HI AS nM CR CY UR UY --readFilesCommand zcat --outFilterMultimapNmax 100 --winAnchorMultimapNmax 100 --outMultimapperOrder Random --runRNGseed 777 --outSAMmultNmax 1’. The default scTE parameters for 10x were used to get the molecule count matrix. The count matrix was lightly filtered to exclude cell barcodes with low numbers of counts: Cells with less than 1000 UMIs and less than 500 genes detected were filtered out, and only the top 10,000 cells with the highest gene count were kept (these default setting can be altered with the ‘--expect-cells, --min_count and --min_genes’ switches in scTE, note that the cell counts are further filtered on a case-by-case basis for each experiment, as detailed below). Other downstream analysis was performed by SCANPY^24^. Specific analysis settings for the individual datasets are described below.

### Analysis of C1/SMART-seq-style data

scRNA-seq data were processed using the scTE C1/SMART–seq pipeline, Briefly, reads were aligned to the genome using STAR^77^, with the setting ‘--winAnchorMultimapNmax 100 --outSAMmultNmax 1 --outSAMmultNmax 1’. The default scTE parameters for C1/SMART-seq were used to get the molecule count matrix. Cells with less than 10,000 counts and less than 2000 expressed genes were filtered out. Cells with more than 20% fraction of mitochondrial counts were discarded. Downstream analysis was performed the same as for the 10x data pipeline. Fluidigm C1/SMART-seq data comes as a single BAM file per barcode. To analyze this data, the ‘barcode’ is taken from the input BAM filenames, and both -CB and -UMI should be False:

~~~
scTE -i *.bam -p 4 -o <output_name> --genome mm10 -x mm10.exclusive.idx -CB False -UMI
False
~~~

The resulting matrices can then be integrated into an scRNA-seq analysis pipeline.

### Analysis of human cardiac differentiation scRNA-seq data

The raw data were download from E-MTAB-6268^33^. As this data was generated using the Single Cell 3’ Library, Gel Bead and Multiplex kit (version 1, 10x Genomics, Cat. #PN-120233), the cell barcode and UMI sequence are not in the same read. First, we merged the cell barcode and UMI sequence into the same read using a custom script, and then aligned the modified fastq file to the hg38 genome using STARsolo, as described above. Cells with less than 500 expressed genes/TEs and cells that have more than 20% fraction of mitochondrial reads were discarded. Single cell trajectory was analyzed by Harmony^78^ and the top 1000 highly variable genes were used for PCA, and the force directed layout was computed using first 150 PCs (principle components). Differentially expressed genes and TEs were analyzed using the SCANPY rank_genes_groups functions by t-test method, the top 500 specifically expressed TEs and genes with Benjamini-Hochberg corrected p-value <0.01 and log2(fold-change) > 0.5 are selected for downstream analysis.

### Analysis of the gastrulation scRNA-seq data

The raw data was download from E-MTAB-6967, and aligned to the mm10 genome using STARsolo^77^, with the parameters ‘--readFilesCommand zcat --outFilterMultimapNmax 100 --winAnchorMultimapNmax 100 --outMultimapperOrder Random --runRNGseed 777 --outSAMmultNmax 1’. Cells with less than 3000 expressed genes/TEs, and less than 8000 UMIs were discarded. Genes expressed in less than 50 cells were removed from the analysis. The count matrix was normalized using normalize_total function of SCANPY, and the top 2000 most highly variable genes were used for PCA, and the first 20 PCs (principle components) were used, as described in the original publication^16^. UMAP plots were generated (min_dist=0.6). Data is from E-MTAB-6967^16^.

### Analysis of Tabula Muris scRNA-seq data

The C1/Smart-seq2 scRNA-seq raw data was download from GSE109774 ^42^, the reads were aligned to the mm10 genome using STAR with the parameters ‘--readFilesCommand zcat --outFilterMultimapNmax 100 --winAnchorMultimapNmax 100 --outMultimapperOrder Random --runRNGseed 777 --outSAMmultNmax 1’. The genes/TEs and cell expression matrix was generated using scTE. Cells with less than 50000 counts or more than 2^7^ counts, less than 1000 expressed genes, or more than 20% fraction of mitochondrial counts were removed. The filtered matrix was normalized using scran^79^. The top 4000 most highly variable genes were used for PCA, and the first 50 PCs were used for downstream analysis. The cell cluster specific expressed genes/TEs was calculated using SCANPY rank_genes_groups functions by t-test method, the top 500 specifically expressed TEs and genes with Benjamini-Hochberg corrected p-value <0.01 and log2(fold-change) >0.5 compare to all other groups of cells were kept.

### Analysis of the OKSM/Chemical reprogramming data

The raw data were download from GSE115943^50^ and GSE114952^29^. Cells with less than 10000 UMIs or more than 1000000 UMIs, or expressed less than 1000 expressed genes, or more than 20% fraction of mitochondrial counts were removed. The filtered matrices were normalized using scran^79^. The top 4000 most highly variable genes were used for PCA, and the first 50 PCs were used for downstream analysis. The cell trajectory routes were taken from the original studies. Differentially expressed genes/TEs were calculated using SCANPY rank_genes_groups functions by the t-test method, the TEs and genes with Benjamini-Hochberg corrected p-value <0.01 and log2(fold-change) >0.5 compared to all other branches of cells were kept.

### Analysis of the OKS reprogramming data

The C1/SMART-seq data were taken from GSE103221 ^25^. the reads were aligned to the mm10 genome using STAR with the parameters ‘--readFilesCommand zcat --outFilterMultimapNmax 100 --winAnchorMultimapNmax 100 --outMultimapperOrder Random --runRNGseed 777 --outSAMmultNmax 1’. The genes/TEs and cells expression matrix was generated using scTE. Cells with less than 10000 counts or more than 2^7^ counts, less than 1000 expressed genes, or more than 20% fraction of mitochondrial counts were removed. The filtered matrix was normalized using scran ^79^. The top 4000 most highly variable genes were used for PCA, and the first 50 PCs were used for downstream analysis. The genes/TEs expression trajectories on pseudotemporal orderings of cells (Fig. 5e) were analyzed by LineagePulse (https://github.com/YosefLab/LineagePulse) according to the pseudotime taken from the original study.

### Analysis of the embryonic heart scRNA-seq data

The raw data was download from GSE126128^40^. This data was aligned to the genome using STARsolo^77^, as described above. Cells with less than 3000 expressed genes/TEs and the cells with less than 8000 UMIs or more than 100000 UMIS were deleted from the analysis. The count matrix was normalized using normalize_total function of SCANPY. The top 2000 most highly variable genes were used for PCA, and the first 20 PCs were used for downstream analysis. UMA projections were generated (min_dist=0.7).

### Analysis of Alzheimer’s disease scRNA-seq data

The MARS-seq scRNA-seq raw data were download from GSE98969 ^65^. The raw fastq file were modified using custom scripts to embed the cell barcode and UMI in the same read, as in the 10x scRNA-seq format. The modified reads were aligned to the mm10 genome with STARsolo as described above. Cells with less than 5000 UMIs or more than 1000000 UMIs, or expressed less than 500 genes, or more than 20% fraction of mitochondrial counts, were removed. The filtered matrix was normalized using scran^79^. The top 4000 most highly variable genes were used for PCA, and the first 50 PCs were used for downstream analysis. The differentially expressed genes and TEs between M2 and M1/3 were analyzed using SCANPY rank_genes_groups functions by t-test method, the genes or TEs with Benjamini-Hochberg corrected p-value <0.01 and log2(fold-change) >0.5 compared to each other were kept.

### Analysis of the Type 2 diabetes/glioblastoma sc-RNA-seq data

The raw data was download from GSE86473^66^, GSE81608^67^. The data was aligned to the hg38 genome using STAR77, as described above for C1 data. Cells with less than 5000 expressed genes/TEs and cells with less than 1*10^6^ counts or more than 6*10^6^ or were deleted from the analysis. The count matrix was normalized using the normalize_total function of SCANPY. There was a strong batch effect based on the sex of the donor in the type 2 diabetes datasets, this was removed using the regress_out function of SCANPY^24^. We did not detect any other batch effect from other confounding variables (age, body-mass index, race). The top 2000 most highly variable genes were used for PCA, and the first 15 PCs (type 2 diabetes) or 25 PCs (glioblastoma) were used. UMAP plots were generated using SCANPY (min_dist=0.7).

### Bulk RNA-seq analysis

Analysis of bulk RNA-seq was performed essentially as previously described^3,80^, with some modifications. Briefly, reads were aligned to the mouse or human genome/transcriptome (GENCODE transcript annotations, mouse M21 or human 30) using STAR (v2.7.1a)^77^. TEtranscripts^81^ or scTE (with the setting -CB False -UMI False) was used to quantitate reads on TEs. Reads were GC normalized using EDASeq (v2.16.3) ^82^, and analyzed using glbase^83^.

### Motif enrichment analysis

The TF motif enrichment in TEs (Supplementary Fig. 6c and 8l) was measured using AME from the MEME suite^84^ with the options “--control --shuffle”.

### Bulk ATAC-seq analysis

Analysis of bulk RNA-seq was performed essentially as previously described^3,85^. Briefly, reads were aligned to the mouse or human genome (mm10 or hg38) using bowtie2 (v2.3.5.1), with the options: “-p 6 --mm --very-sensitive --no-unal --no-mixed --no-discordant -X2000”, and reads mapping to TEs were counted using te_counter (https://github.com/oaxiom/te_counter). The counts per million (CPM) reads metric was used for enrichment scores.

### Analysis of the scATAC-seq data

We downloaded the scATAC-seq data from the 10x Illumina website (https://support.10xgenomics.com/single-cell-atac/datasets/1.1.0/atac_pbmc_10k_v1).The barcode was inserted into the read name, so that the mapping could keep track of the cell ID. This yielded reads names inside the FASTQ, such as: (where CCACGTTGTGGACTGA sequence is the cell barcode)

~~~
@CCACGTTGTGGACTGA:A00519:269:H7FM2DRXX:1:1101:1325:1000 1:N:0:AAGCATAA
~~~

The data was aligned to the human hg38 genome using bowtie2 ^86^ with the command options “-p 6 --mm --very-sensitive --no-unal --no-mixed --no-discordant -X2000”. The resulting data was then processed using scTE with the command:

~~~
scTE_scatacseq -i $<in> -x hg38.te.atac.idx -g hg38 -p 1 -UMI False -CB True -o <out>
~~~

The genome indices were prebuilt using:

~~~
wget -c -O mm10.te.txt.gz ‘http://hgdownload.soe.ucsc.edu/goldenPath/mm10/database/rmsk.txt.gz’
zcat mm10.te.txt.gz | grep -E ‘LINE|SINE|LTR|Retroposon’ | cut -f6-8,11 >mm10.te.bed
python3 /share/apps/genomics/unstable/scTE/bin/scTEATAC_build -g mm10.te.bed -o mm10.te.atac
wget -c -O hg38.te.txt.gz ‘http://hgdownload.soe.ucsc.edu/goldenPath/hg38/database/rmsk.txt.gz’
zcat hg38.te.txt.gz | grep -E ‘LINE|SINE|LTR|Retroposon’ | cut -f6-8,11 >hg38.te.bed
python3 /share/apps/genomics/unstable/scTE/bin/scTEATAC_build -g hg38.te.bed -o hg38.te.atac
~~~

## Acknowledgements

We grateful to Rujin Huang for the helpful discussions and advice. We appreciate the assistance of Lihui Lin, Huijian Feng and Yuanbang Mai on the data analysis. We thank the GuangZhou Branch of the Supercomputing Center of Chinese Academy of Science for its support. We thank the support from the Center for Computational Science and Engineering of Southern University of Science and Technology. This work was supported by the National Natural Science Foundation of China (31970589, 31801217, 31850410463, 31850410486), Guangdong Science and Technology Commission (2019A050510004), and the Shenzhen Peacock plan (201701090668B).

## Author contributions

J.H., A.P.H., and J.C. initiated the project and wrote the manuscript. J.H. and A.P.H. performed the bioinformatic analysis with the assistance of all other authors. A.P.H and J.C. supervised and funded the project.

## Competing interests

The authors declare no competing interests

## Notes

### Competing Interest Statement

The authors have declared no competing interest.

## References

1 Cordaux, R. & Batzer, M. A. The impact of retrotransposons on human genome evolution. Nat Rev Genet 10, 691–703, doi:10.1038/nrg2640 (2009).

2 Hutchins, A. P. & Pei, D. Transposable elements at the center of the crossroads between embryogenesis, embryonic stem cells, reprogramming, and long non-coding RNAs. Sci Bull 60, 1722–1733, doi:10.1007/s11434-015-0905-x (2015).

3 He, J. et al. Transposable elements are regulated by context-specific patterns of chromatin marks in mouse embryonic stem cells. Nat Commun 10, 34, doi:10.1038/s41467-018-08006-y (2019).

4 Lu, X. et al. The retrovirus HERVH is a long noncoding RNA required for human embryonic stem cell identity. Nat Struct Mol Biol 21, 423–425, doi:10.1038/nsmb.2799 (2014).

5 Bourque, G. et al. Ten things you should know about transposable elements. Genome Biol 19, 199, doi:10.1186/s13059-018-1577-z (2018).

6 Goke, J. et al. Dynamic transcription of distinct classes of endogenous retroviral elements marks specific populations of early human embryonic cells. Cell Stem Cell 16, 135–141, doi:10.1016/j.stem.2015.01.005 (2015).

7 Grow, E. J. et al. Intrinsic retroviral reactivation in human preimplantation embryos and pluripotent cells. Nature 522, 221–225, doi:10.1038/nature14308 (2015).

8 Percharde, M. et al. A LINE1-Nucleolin Partnership Regulates Early Development and ESC Identity. Cell 174, 391-405.e319, doi:10.1016/j.cell.2018.05.043 (2018).

9 Jachowicz, J. W. et al. LINE-1 activation after fertilization regulates global chromatin accessibility in the early mouse embryo. Nat Genet 49, 1502–1510, doi:10.1038/ng.3945 (2017).

10 Tam, O. H., Ostrow, L. W. & Gale Hammell, M. Diseases of the nERVous system: retrotransposon activity in neurodegenerative disease. Mobile DNA 10, 32, doi:10.1186/s13100-019-0176-1 (2019).

11 Jang, H. S. et al. Transposable elements drive widespread expression of oncogenes in human cancers. Nat Genet 51, 611–617, doi:10.1038/s41588-019-0373-3 (2019).

12 Chuong, E. B., Elde, N. C. & Feschotte, C. Regulatory activities of transposable elements: from conflicts to benefits. Nat Rev Genet 18, 71–86, doi:10.1038/nrg.2016.139 (2017).

13 Feschotte, C. Transposable elements and the evolution of regulatory networks. Nat Rev Genet 9, 397–405, doi:10.1038/nrg2337 (2008).

14 Kunarso, G. et al. Transposable elements have rewired the core regulatory network of human embryonic stem cells. Nat Genet 42, 631–634, doi:10.1038/ng.600 (2010).

15 Faulkner, G. J. et al. The regulated retrotransposon transcriptome of mammalian cells. Nat Genet 41, 563–571, doi:10.1038/ng.368 (2009).

16 Pijuan-Sala, B. et al. A single-cell molecular map of mouse gastrulation and early organogenesis. Nature 566, 490–495, doi:10.1038/s41586-019-0933-9 (2019).

17 Li, H. et al. Dysfunctional CD8 T Cells Form a Proliferative, Dynamically Regulated Compartment within Human Melanoma. Cell 176, 775–789 e718, doi:10.1016/j.cell.2018.11.043 (2019).

18 Tirosh, I. et al. Dissecting the multicellular ecosystem of metastatic melanoma by single-cell RNA-seq. Science (New York, N.Y.) 352, 189–196, doi:10.1126/science.aad0501 (2016).

19 Macfarlan, T. S. et al. Embryonic stem cell potency fluctuates with endogenous retrovirus activity. Nature 487, 57–63, doi:10.1038/nature11244 (2012).

20 Fu, X., Wu, X., Djekidel, M. N. & Zhang, Y. Myc and Dnmt1 impede the pluripotent to totipotent state transition in embryonic stem cells. Nat Cell Biol 21, 835–844, doi:10.1038/s41556-019-0343-0 (2019).

21 Treangen, T. J. & Salzberg, S. L. Repetitive DNA and next-generation sequencing: computational challenges and solutions. Nat Rev Genet 13, 36–46, doi:10.1038/nrg3117 (2011).

22 Goerner-Potvin, P. & Bourque, G. Computational tools to unmask transposable elements. Nat Rev Genet 19, 688–704,,doi:10.1038/s41576-018-0050-x (2018).

23 Stuart, T. et al. Comprehensive Integration of Single-Cell Data. Cell 177, 1888–1902 e1821, doi:10.1016/j.cell.2019.05.031 (2019).

24 Wolf, F. A., Angerer, P. & Theis, F. J. SCANPY: large-scale single-cell gene expression data analysis. Genome Biol 19, 15, doi:10.1186/s13059-017-1382-0 (2018).

25 Guo, L. et al. Resolving Cell Fate Decisions during Somatic Cell Reprogramming by Single-Cell RNA-Seq. Mol Cell 73, 815–829 e817, doi:10.1016/j.molcel.2019.01.042 (2019).

26 Zheng, G. X. et al. Massively parallel digital transcriptional profiling of single cells. Nat Commun 8, 14049, doi:10.1038/ncomms14049 (2017).

27 Garcia-Perez, J. L., Widmann, T. J. & Adams, I. R. The impact of transposable elements on mammalian development. Development 143, 4101–4114, doi:10.1242/dev.132639 (2016).

28 Rodriguez-Terrones, D. & Torres-Padilla, M. E. Nimble and Ready to Mingle: Transposon Outbursts of Early Development. Trends in genetics: TIG 34, 806–820, doi:10.1016/j.tig.2018.06.006 (2018).

29 Zhao, T. et al. Single-Cell RNA-Seq Reveals Dynamic Early Embryonic-like Programs during Chemical Reprogramming. Cell Stem Cell 23, 31–45 e37, doi:10.1016/j.stem.2018.05.025 (2018).

30 Wang, J. et al. Primate-specific endogenous retrovirus-driven transcription defines naive-like stem cells. Nature 516, 405–409, doi:10.1038/nature13804 (2014).

31 Fort, A. et al. Deep transcriptome profiling of mammalian stem cells supports a regulatory role for retrotransposons in pluripotency maintenance. Nat Genet 46, 558–566, doi:10.1038/ng.2965 (2014).

32 Theunissen, T. W. et al. Molecular Criteria for Defining the Naive Human Pluripotent State. Cell Stem Cell 19, 502–515, doi:10.1016/j.stem.2016.06.011 (2016).

33 Friedman, C. E. et al. Single-Cell Transcriptomic Analysis of Cardiac Differentiation from Human PSCs Reveals HOPX-Dependent Cardiomyocyte Maturation. Cell Stem Cell 23, 586–598 e588, doi:10.1016/j.stem.2018.09.009 (2018).

34 Liu, Q. et al. Genome-Wide Temporal Profiling of Transcriptome and Open Chromatin of Early Cardiomyocyte Differentiation Derived From hiPSCs and hESCs. Circ Res 121, 376–391, doi:10.1161/CIRCRESAHA.116.310456 (2017).

35 Abed, M. et al. The Gag protein PEG10 binds to RNA and regulates trophoblast stem cell lineage specification. PLoS One 14, e0214110, doi:10.1371/journal.pone.0214110 (2019).

36 Zhong, Y. et al. Isolation of primitive mouse extraembryonic endoderm (pXEN) stem cell lines. Stem Cell Res 30, 100–112, doi:10.1016/j.scr.2018.05.008 (2018).

37 Factor, D. C. et al. Epigenomic comparison reveals activation of “seed” enhancers during transition from naive to primed pluripotency. Cell Stem Cell 14, 854–863, doi:10.1016/j.stem.2014.05.005 (2014).

38 Morey, L. et al. Polycomb Regulates Mesoderm Cell Fate-Specification in Embryonic Stem Cells through Activation and Repression Mechanisms. Cell Stem Cell 17, 300–315, doi:10.1016/j.stem.2015.08.009 (2015).

39 Merkin, J., Russell, C., Chen, P. & Burge, C. B. Evolutionary dynamics of gene and isoform regulation in Mammalian tissues. Science (New York, N.Y.) 338, 1593–1599, doi:10.1126/science.1228186 (2012).

40 de Soysa, T. Y. et al. Single-cell analysis of cardiogenesis reveals basis for organ-level developmental defects. Nature 572, 120–124, doi:10.1038/s41586-019-1414-x (2019).

41 Richardson, S. R., Morell, S. & Faulkner, G. J. L1 retrotransposons and somatic mosaicism in the brain. Annu Rev Genet 48, 1–27, doi:10.1146/annurev-genet-120213-092412 (2014).

42 Tabula Muris, C. et al. Single-cell transcriptomics of 20 mouse organs creates a Tabula Muris. Nature 562, 367–372, doi:10.1038/s41586-018-0590-4 (2018).

43 Chuong, E. B., Elde, N. C. & Feschotte, C. Regulatory evolution of innate immunity through co-option of endogenous retroviruses. Science (New York, N.Y.) 351, 1083–1087, doi:10.1126/science.aad5497 (2016).

44 Koonin, E. V. & Krupovic, M. Evolution of adaptive immunity from transposable elements combined with innate immune systems. Nat Rev Genet 16, 184–192, doi:10.1038/nrg3859 (2015).

45 Erwin, J. A., Marchetto, M. C. & Gage, F. H. Mobile DNA elements in the generation of diversity and complexity in the brain. Nat Rev Neurosci 15, 497–506, doi:10.1038/nrn3730 (2014).

46 Takahashi, K. & Yamanaka, S. Induction of pluripotent stem cells from mouse embryonic and adult fibroblast cultures by defined factors. Cell 126, 663–676, doi:10.1016/j.cell.2006.07.024 (2006).

47 Wang, B. et al. Induction of Pluripotent Stem Cells from Mouse Embryonic Fibroblasts by Jdp2-Jhdm1b-Mkk6-Glis1-Nanog-Essrb-Sall4. Cell Rep 27, 3473–3485 e3475, doi:10.1016/j.celrep.2019.05.068 (2019).

48 Hou, P. et al. Pluripotent stem cells induced from mouse somatic cells by small-molecule compounds. Science (New York, N.Y.) 341, 651–654, doi:10.1126/science.1239278 (2013).

49 Cao, S. et al. Chromatin Accessibility Dynamics during Chemical Induction of Pluripotency. Cell Stem Cell 22, 529–542 e525, doi:10.1016/j.stem.2018.03.005 (2018).

50 Schiebinger, G. et al. Optimal-Transport Analysis of Single-Cell Gene Expression Identifies Developmental Trajectories in Reprogramming. Cell 176, 928–943 e922, doi:10.1016/j.cell.2019.01.006 (2019).

51 Friedli, M. et al. Loss of transcriptional control over endogenous retroelements during reprogramming to pluripotency. Genome Res 24, 1251–1259, doi:10.1101/gr.172809.114 (2014).

52 Liu, J. et al. The oncogene c-Jun impedes somatic cell reprogramming. Nat Cell Biol 17, 856–867, doi:10.1038/ncb3193 (2015).

53 Chronis, C. et al. Cooperative Binding of Transcription Factors Orchestrates Reprogramming. Cell 168, 442–459 e420, doi:10.1016/j.cell.2016.12.016 (2017).

54 Buenrostro, J. D. et al. Single-cell chromatin accessibility reveals principles of regulatory variation. Nature 523, 486–490, doi:10.1038/nature14590 (2015).

55 Shema, E., Bernstein, B. E. & Buenrostro, J. D. Single-cell and single-molecule epigenomics to uncover genome regulation at unprecedented resolution. Nat Genet 51, 19–25, doi:10.1038/s41588-018-0290-x (2019).

56 Cusanovich, D. A. et al. A Single-Cell Atlas of In Vivo Mammalian Chromatin Accessibility. Cell 174, 1309–1324 e1318, doi:10.1016/j.cell.2018.06.052 (2018).

57 Sun, X. et al. Transcription factor profiling reveals molecular choreography and key regulators of human retrotransposon expression. Proc Natl Acad Sci U S A 115, E5526–E5535, doi:10.1073/pnas.1722565115 (2018).

58 Chen, X., Miragaia, R. J., Natarajan, K. N. & Teichmann, S. A. A rapid and robust method for single cell chromatin accessibility profiling. Nat Commun 9, 5345, doi:10.1038/s41467-018-07771-0 (2018).

59 Warren, J. S. et al. Histone methyltransferase Smyd1 regulates mitochondrial energetics in the heart. Proc Natl Acad Sci U S A 115, E7871–E7880, doi:10.1073/pnas.1800680115 (2018).

60 Rasmussen, T. L. et al. Smyd1 facilitates heart development by antagonizing oxidative and ER stress responses. PLoS One 10, e0121765, doi:10.1371/journal.pone.0121765 (2015).

61 Franklin, S. et al. The chromatin-binding protein Smyd1 restricts adult mammalian heart growth. Am J Physiol Heart Circ Physiol 311, H1234–H1247, doi:10.1152/ajpheart.00235.2016 (2016).

62 Calderon, D. et al. Landscape of stimulation-responsive chromatin across diverse human immune cells. Nat Genet, doi:10.1038/s41588-019-0505-9 (2019).

63 Bravo Gonzalez-Blas, C. et al. cisTopic: cis-regulatory topic modeling on single-cell ATAC-seq data. Nat Methods 16, 397–400, doi:10.1038/s41592-019-0367-1 (2019).

64 Li, W. et al. Activation of transposable elements during aging and neuronal decline in Drosophila. Nat Neurosci 16, 529–531, doi:10.1038/nn.3368 (2013).

65 Keren-Shaul, H. et al. A Unique Microglia Type Associated with Restricting Development of Alzheimer’s Disease. Cell 169, 1276–1290 e1217, doi:10.1016/j.cell.2017.05.018 (2017).

66 Lawlor, N. et al. Single-cell transcriptomes identify human islet cell signatures and reveal cell-type-specific expression changes in type 2 diabetes. Genome Res 27, 208–222, doi:10.1101/gr.212720.116 (2017).

67 Xin, Y. et al. RNA Sequencing of Single Human Islet Cells Reveals Type 2 Diabetes Genes. Cell Metab 24, 608–615, doi:10.1016/j.cmet.2016.08.018 (2016).

68 Fadista, J. et al. Global genomic and transcriptomic analysis of human pancreatic islets reveals novel genes influencing glucose metabolism. Proc Natl Acad Sci U S A 111, 13924–13929, doi:10.1073/pnas.1402665111 (2014).

69 Darmanis, S. et al. Single-Cell RNA-Seq Analysis of Infiltrating Neoplastic Cells at the Migrating Front of Human Glioblastoma. Cell Rep 21, 1399–1410, doi:10.1016/j.celrep.2017.10.030 (2017).

70 Chuong, E. B., Rumi, M. A., Soares, M. J. & Baker, J. C. Endogenous retroviruses function as species-specific enhancer elements in the placenta. Nat Genet 45, 325–329, doi:10.1038/ng.2553 (2013).

71 Payer, L. M. & Burns, K. H. Transposable elements in human genetic disease. Nat Rev Genet, doi:10.1038/s41576-019-0165-8 (2019).

72 Burns, K. H. Transposable elements in cancer. Nat Rev Cancer 17, 415–424, doi:10.1038/nrc.2017.35 (2017).

73 De Cecco, M. et al. L1 drives IFN in senescent cells and promotes age-associated inflammation. Nature 566, 73–78, doi:10.1038/s41586-018-0784-9 (2019).

74 Li, H. et al. The Sequence Alignment/Map format and SAMtools. Bioinformatics 25, 2078–2079, doi:10.1093/bioinformatics/btp352 (2009).

75 Frankish, A. et al. GENCODE reference annotation for the human and mouse genomes. Nucleic Acids Res 47, D766–D773, doi:10.1093/nar/gky955 (2019).

76 Jurka, J. Repbase update: a database and an electronic journal of repetitive elements. Trends in genetics: TIG 16, 418–420, doi:10.1016/s0168-9525(00)02093-x (2000).

77 Dobin, A. et al. STAR: ultrafast universal RNA-seq aligner. Bioinformatics 29, 15–21, doi:10.1093/bioinformatics/bts635 (2013).

78 Nowotschin, S. et al. The emergent landscape of the mouse gut endoderm at single-cell resolution. Nature 569, 361–367, doi:10.1038/s41586-019-1127-1 (2019).

79 Lun, A. T., McCarthy, D. J. & Marioni, J. C. A step-by-step workflow for low-level analysis of single-cell RNA-seq data with Bioconductor. F1000Res 5, 2122, doi:10.12688/f1000research.9501.2 (2016).

80 Hutchins, A. P. et al. Models of global gene expression define major domains of cell type and tissue identity. Nucleic Acids Res 45, 2354–2367, doi:10.1093/nar/gkx054 (2017).

81 Jin, Y., Tam, O. H., Paniagua, E. & Hammell, M. TEtranscripts: a package for including transposable elements in differential expression analysis of RNA-seq datasets. Bioinformatics 31, 3593–3599, doi:10.1093/bioinformatics/btv422 (2015).

82 Risso, D., Schwartz, K., Sherlock, G. & Dudoit, S. GC-content normalization for RNA-Seq data. BMC Bioinformatics 12, 480, doi:10.1186/1471-2105-12-480 (2011).

83 Hutchins, A. P., Jauch, R., Dyla, M. & Miranda-Saavedra, D. glbase: a framework for combining, analyzing and displaying heterogeneous genomic and high-throughput sequencing data. Cell Regen (Lond) 3, 1, doi:10.1186/2045-9769-3-1 (2014).

84 McLeay, R. C. & Bailey, T. L. Motif Enrichment Analysis: a unified framework and an evaluation on ChIP data. BMC Bioinformatics 11, 165, doi:10.1186/1471-2105-11-165 (2010).

85 Li, D. et al. Chromatin Accessibility Dynamics during iPSC Reprogramming. Cell Stem Cell 21, 819–833 e816, doi:10.1016/j.stem.2017.10.012 (2017).

86 Langmead, B. & Salzberg, S. L. Fast gapped-read alignment with Bowtie 2. Nat Methods 9, 357–359, doi:10.1038/nmeth.1923 (2012).

