## Supplemental figures for "Unveiling transposable element expression heterogeneity in cell fate regulation at the single-cell level"

### fig. S1

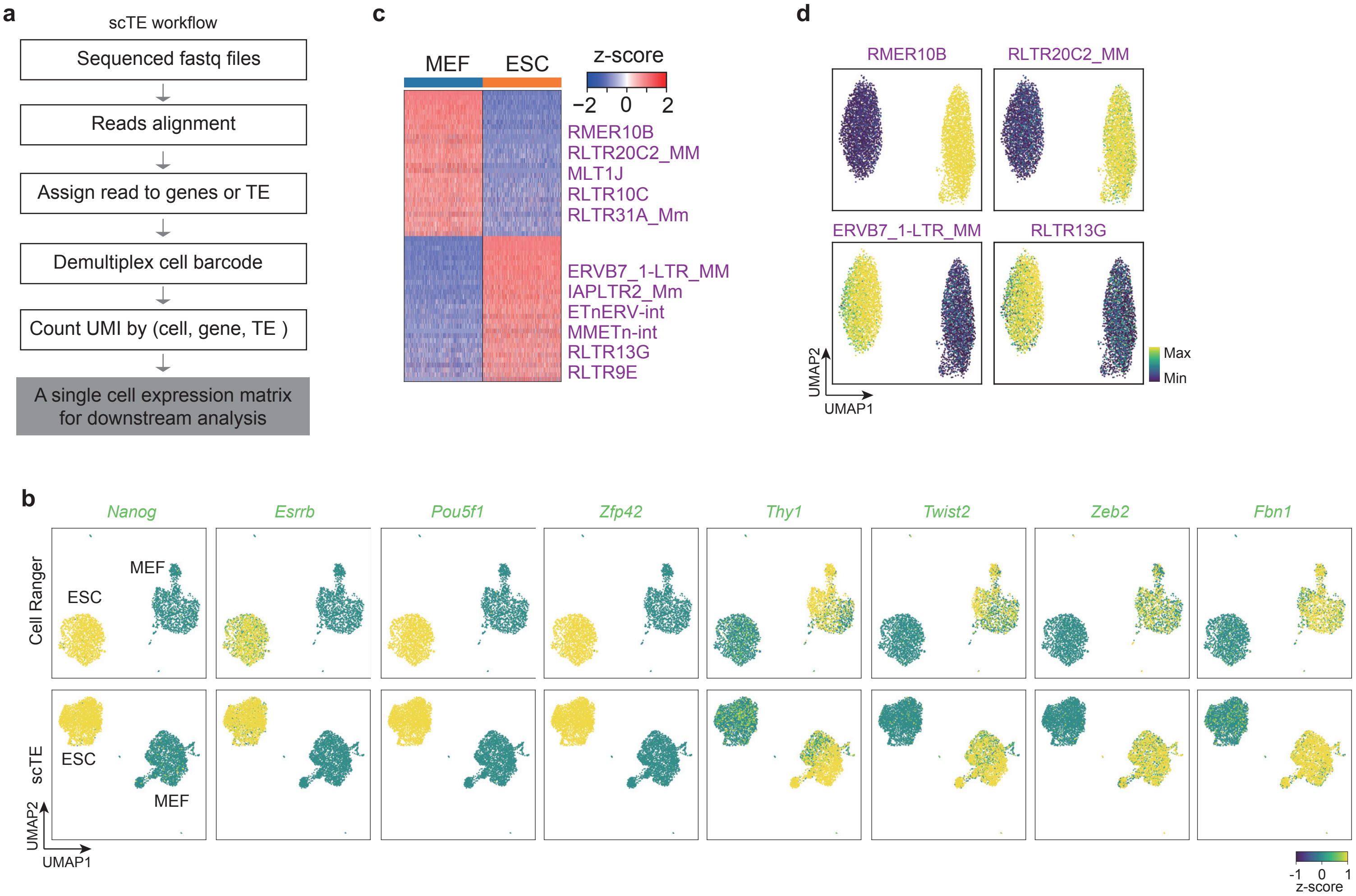
a
scTE workflow
Sequenced fastq files
Reads alignment
Assign read to genes or TE
Demultiplex cell barcode
Count UMI by (cell, gene, TE )
A single cell expression matrix for downstream analysis
c
MEF
ESC
z-score
−2 0 2
RMER10B
RLTR20C2_MM
MLT1J
RLTR10C
RLTR31A_Mm
ERVB7_1-LTR_MM
IAPLTR2_Mm
ETnERV-int
MMETn-int
RLTR13G
RLTR9E
d
RMER10B
RLTR20C2_MM
ERVB7_1-LTR_MM
RLTR13G
Max
Min
UMAP2
UMAP1
b
Nanog
Esrrb
Pou5f1
Zfp42
Thy1
Twist2
Zeb2
Fbn1
Cell Ranger
scTE
ESC
MEF
UMAP2
UMAP1
-1 0 1
z-score

**Supplementary Fig. 1 l Comparison of scTE versus Cell Ranger analysis.** (**a**) Schematic describing the scTE workflow. (**b**) As Fig. 1b, but cells are colored by the expression of indicated genes. (**c**) Heatmap of TE expression differences between MEF and ESC single cells. Selected differentially expressed TEs are labelled. (**d**) Expression of selected TEs in a 50:50 split of MEF and ESC data.

fig S2

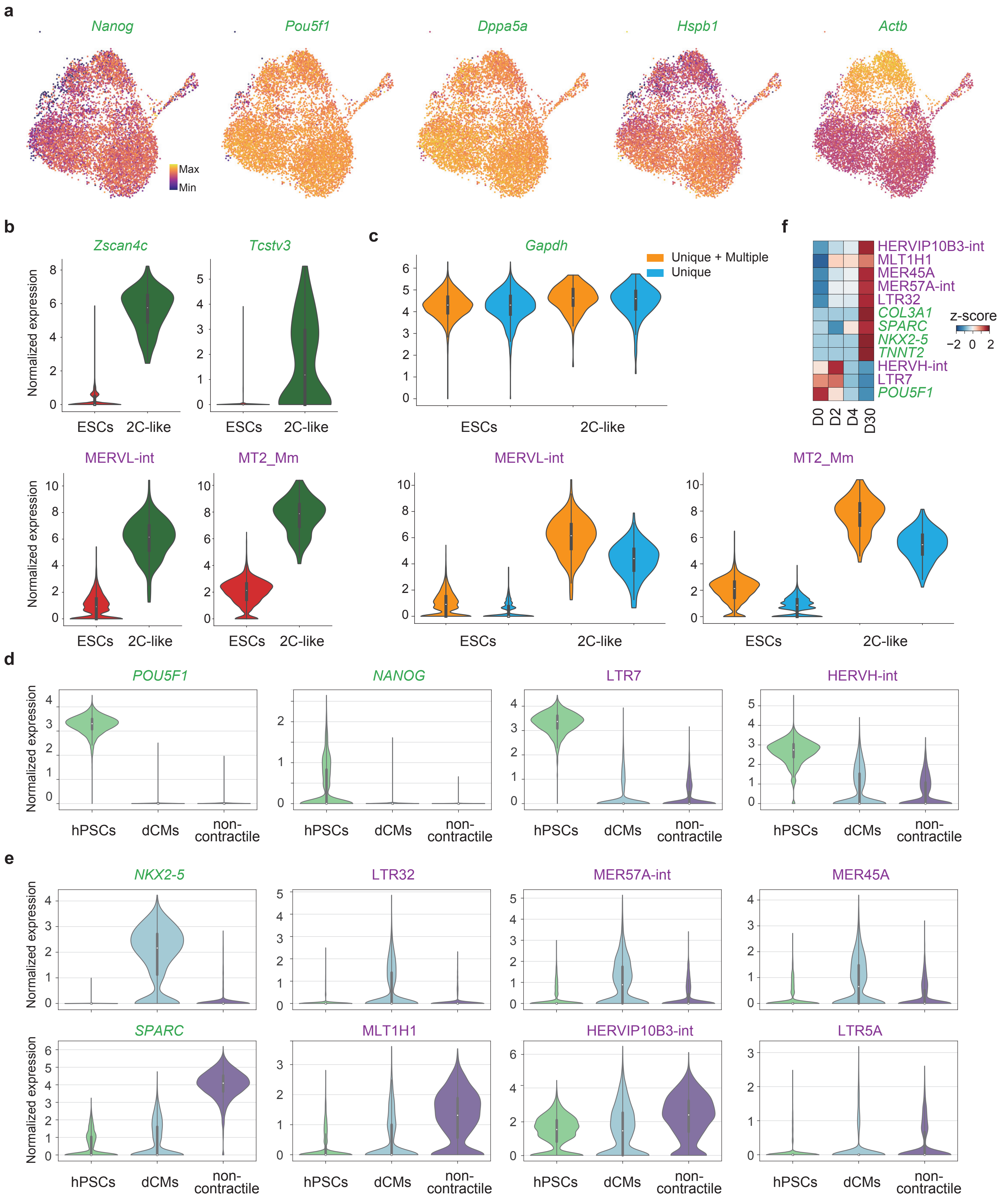

a
Nanog
Pou5f1
Dppa5a
Hspb1
Actb
Max
Min
b
Zscan4c
Tcstv3
Normalized expression
ESCs
2C-like
MERVL-int
MT2_Mm
c
Gapdh
Unique + Multiple
Unique
f
HERVIP10B3-int
MLT1H1
MER45A
MER57A-int
LTR32
COL3A1
SPARC
NKX2-5
TNNT2
HERVH-int
LTR7
POU5F1
z-score
D0
D2
D4
D30
d
POU5F1
NANOG
LTR7
HERVH-int
hPSCs
dCMs
non-contractile
e
NKX2-5
LTR32
MER57A-int
MER45A
SPARC
MLT1H1
HERVIP10B3-int
LTR5A

**Supplementary Fig. 2 I Dynamic expression of TEs in ESCs, and in cardiac differentiation.** (**a**) UMAP plots for mouse ESCs, are colored by the expression of indicated genes. (**b**) Violin plots showing the expression of 2C maker genes/TEs. (**c**) Violin plots showing the expression of indicated gene/TEs with all mapped reads (unique + multiple) or only the unique mapped reads (unique). (**d**) Violin plots showing the expression of a selection of differentially expressed genes and TEs in hPSCs. (**e**) Violin plots showing the expression of a selection of genes specific to the dCM or non-contractile branch cells. (**f**) Expression dynamics of selected TEs and marker genes during cardiac differentiation analyzed using bulk RNA-seq. D indicates the day of the differentiation process.

### fig S3

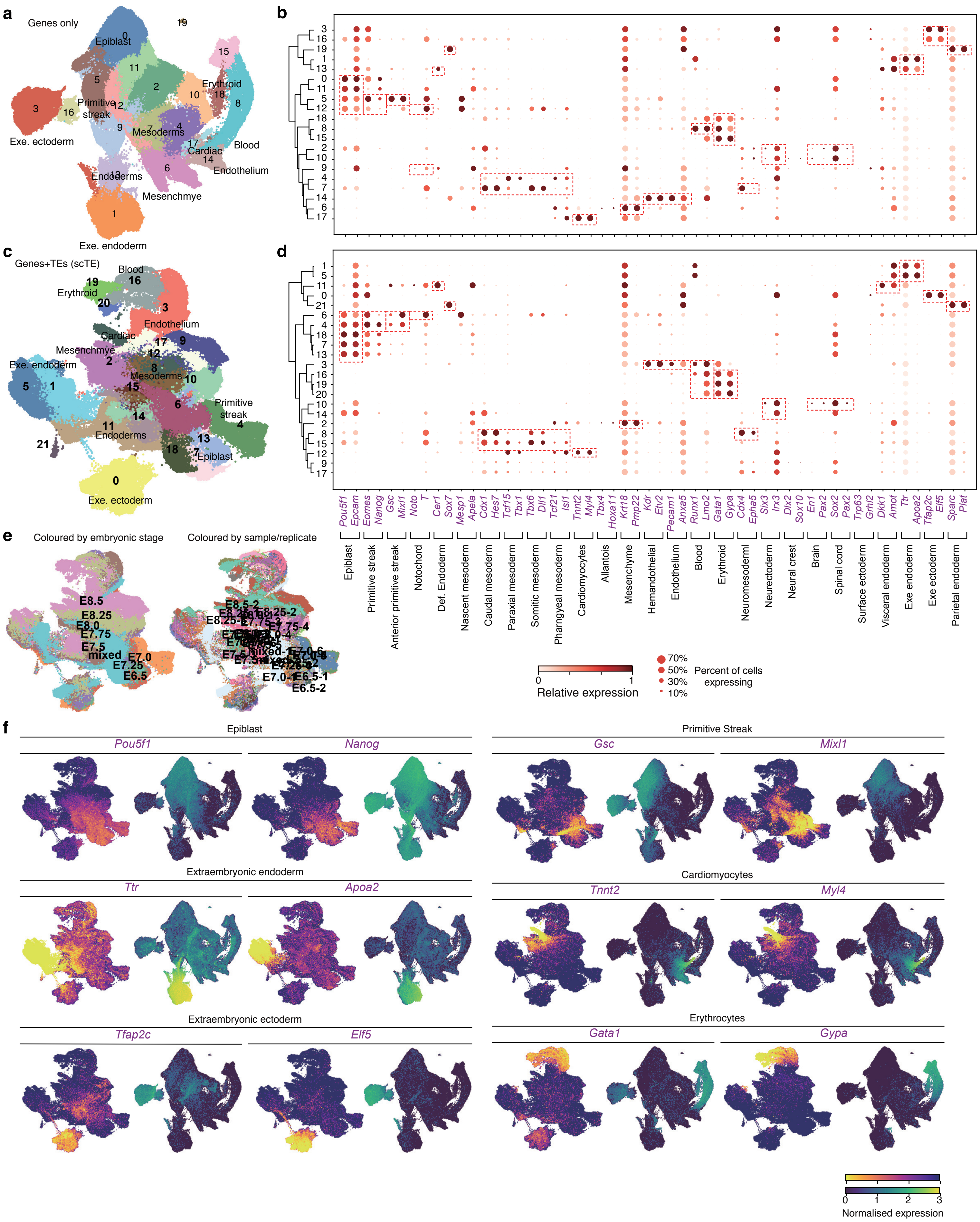

a
Genes only
Epiblast
Primitive streak
Erythroid
Mesoderms
Cardiac
Blood
Endothelium
Endoderms
Mesenchmye
Exe. ectoderm
Exe. endoderm
b
c
Genes+TEs (scTE)
Blood
Erythroid
Endothelium
Cardiac
Mesenchmye
Exe. endoderm
Mesoderms
Primitive streak
Endoderms
Epiblast
Exe. ectoderm
d
Pou5f1 Epcam Eomes Nanog Gsc Mixl1 Noto T Cer1 Sox7 Mesp1 Apela Cdx1 Hes7 Tcf15 Tbx1 Tbx6 Dll1 Tcf21 Isl1 Tnnt2 Myl4 Tbx4 Hoxa11 Krt18 Pmp22 Kdr Etv2 Pecam1 Anxa5 Runx1 Lmo2 Gata1 Gypa Cdx4 Epha5 Six3 Irx3 Dlx2 Sox10 En1 Pax2 Sox2 Pax2 Trp63 Grhl2 Dkk1 Amot Ttr Apoa2 Tfap2c Elf5 Sparc Plat
Epiblast
Primitive streak
Anterior primitive streak
Notochord
Def. Endoderm
Nascent mesoderm
Caudal mesoderm
Paraxial mesoderm
Somitic mesoderm
Pharngyeal mesoderm
Cardiomyocytes
Allantois
Mesenchyme
Hemandothelial
Endothelium
Blood
Erythroid
Neuromesoderml
Neurectoderm
Neural crest
Brain
Spinal cord
Surface ectoderm
Visceral endoderm
Exe endoderm
Exe ectoderm
Parietal endoderm
Relative expression
Percent of cells expressing
70%
50%
30%
10%
e
Coloured by embryonic stage
Coloured by sample/replicate
E8.5
E8.25
E8.0
E7.75
E7.5
mixed
E7.0
E7.25
E6.5
E6.5-1
E6.5-2
f
Epiblast
Pou5f1
Nanog
Primitive Streak
Gsc
Mixl1
Extraembryonic endoderm
Ttr
Apoa2
Cardiomyocytes
Tnnt2
Myl4
Extraembryonic ectoderm
Tfap2c
Elf5
Erythrocytes
Gata1
Gypa
Normalised expression

**Supplementary Fig. 3 I Comparison of gene-based analysis to the gene/TE-based analysis from scTE**. (**a**) Mouse gastrulation was reanalyzed using a gene-based pipeline with STARsolo. Shown are the UMAP plots, labelled with clusters using the Leiden algorithm (resolution=0.5). Selected lineages are indicated. (**b**) Dot plots showing the expression level and the indicated percent of cells the gene is expressed in, for the indicated marker genes. This panel shares the x-axis with panel d. (**c**) Analysis of the gastrulation data using scTE. Cells were plotted by UMAP, with Leiden grouping (resolution=0.5). Selected lineages are labelled. (**d**) Dot plot showing the expression of selected marker genes, as in panel b. Panels b and d share axes and legends. (**e**) UMAP plot of the scTE-analyzed gastrulation data, colored by the embryonic stage, or by the sample/replicate. (**f**) A selection of UMAP plots showing the expression of the indicated marker genes in the gene+TE-based analysis (left), or the gene-based analysis (right).

fig S4

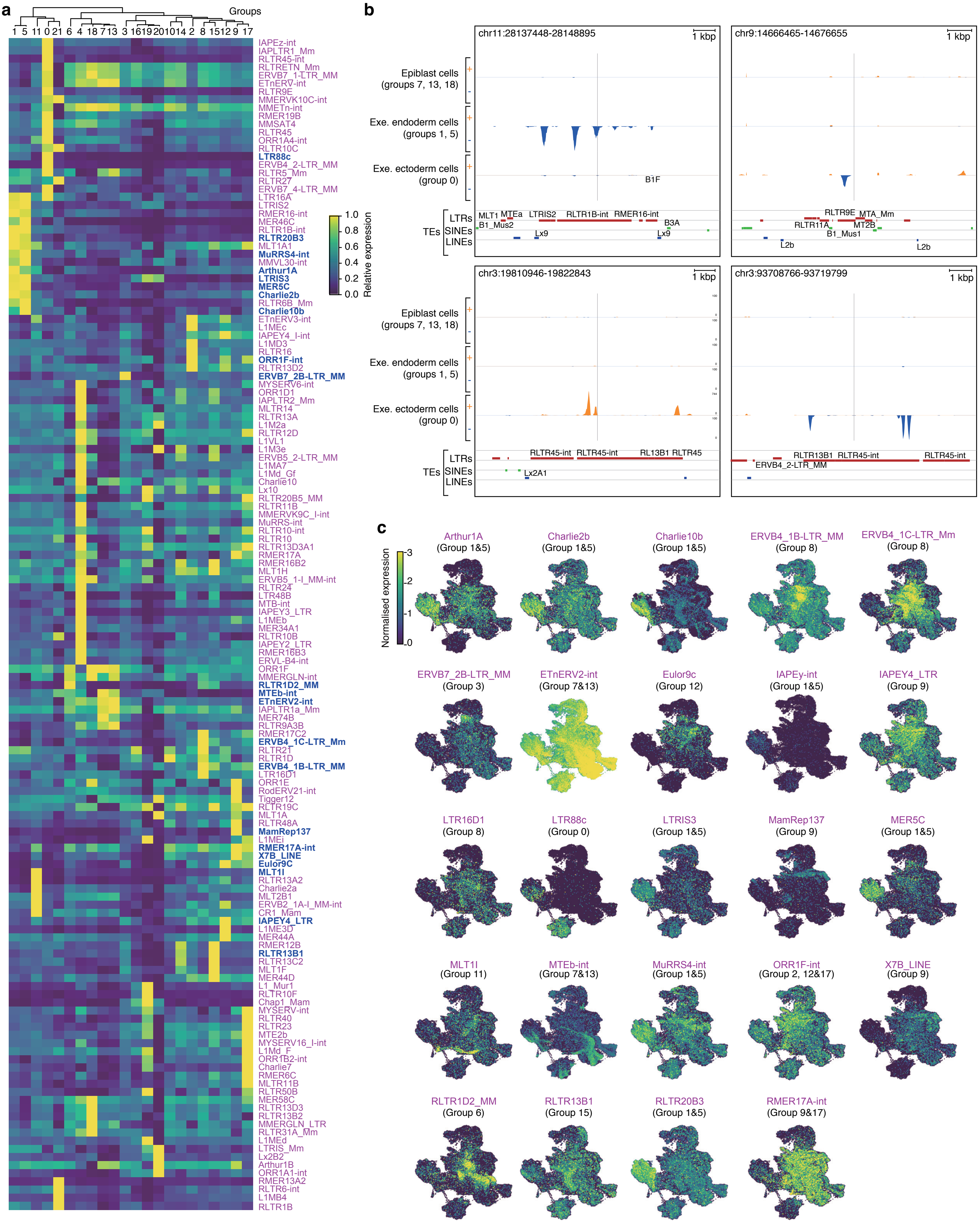
a
Groups
1 5 11 0 21 6 4 18 7 13 3 16 19 20 10 14 2 8 15 12 9 17
Relative expression
b
chr11:28137448-28148895
chr9:14666465-14676655
chr3:19810946-19822843
chr3:93708766-93719799
1 kbp
Epiblast cells (groups 7, 13, 18)
Exe. endoderm cells (groups 1, 5)
Exe. ectoderm cells (group 0)
TEs
LTRs
SINEs
LINEs
c
Normalised expression
Arthur1A (Group 1&5)
Charlie2b (Group 1&5)
Charlie10b (Group 1&5)
ERVB4_1B-LTR_MM (Group 8)
ERVB4_1C-LTR_Mm (Group 8)
ERVB7_2B-LTR_MM (Group 3)
ETnERV2-int (Group 7&13)
Eulor9c (Group 12)
IAPEy-int (Group 1&5)
IAPEY4_LTR (Group 9)
LTR16D1 (Group 8)
LTR88c (Group 0)
LTRIS3 (Group 1&5)
MamRep137 (Group 9)
MER5C (Group 1&5)
MLT1I (Group 11)
MTEb-int (Group 7&13)
MuRRS4-int (Group 1&5)
ORR1F-int (Group 2, 12&17)
X7B_LINE (Group 9)
RLTR1D2_MM (Group 6)
RLTR13B1 (Group 15)
RLTR20B3 (Group 1&5)
RMER17A-int (Group 9&17)

**Supplementary Fig. 4 l Expression of TEs in the gastrulation data.** (**a**) Heatmap showing the expression level of the indicated TEs in the groups, as defined in Fig. S3C. Expression was transformed into the unit variance, and the groups were clustered. The TEs shown here were significantly different (Benjamini-Hochberg corrected Wilcoxon rank-sum test, p-value<0.01), and at least >2-fold change between groups. (**b**) Genome views showing example individual loci. The reads from the gastrulation scRNA-seq data were split according to the groups corresponding to the epiblast, extraembryonic endoderm, or extra embryonic ectoderm cells, as defined in Supplementary Fig. 3c. Shown here are selected genome views indicating the 3' expression of extraembryonic endoderm, or ectoderm-specific TEs.'+' and '-' indicates the strand. Care should be taken in the interpretation of these genome views, and we show them for illustrative purposes only, to show typical read distribution from the 10x data. (**c**) UMAP plots for a selection of TEs, as labelled in blue in panel a.

### fig S5

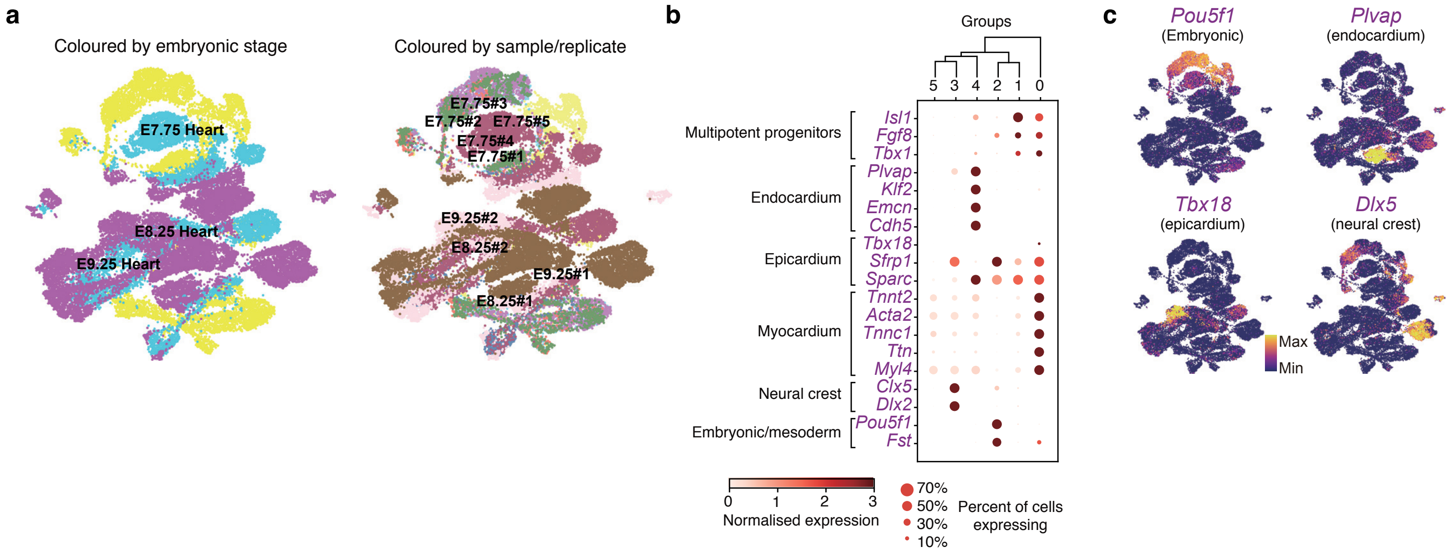

a
Coloured by embryonic stage
E7.75 Heart
E8.25 Heart
E9.25 Heart
Coloured by sample/replicate
E7.75#3
E7.75#2
E7.75#5
E7.75#4
E7.75#1
E9.25#2
E8.25#2
E9.25#1
E8.25#1
b
Groups
5 3 4 2 1 0
Multipotent progenitors
Isl1
Fgf8
Tbx1
Endocardium
Plvap
Klf2
Emcn
Cdh5
Epicardium
Tbx18
Sfrp1
Sparc
Myocardium
Tnnt2
Acta2
Tnnc1
Ttn
Myl4
Neural crest
Clx5
Dlx2
Embryonic/mesoderm
Pou5f1
Fst
0 1 2 3
Normalised expression
70%
50%
30%
10%
Percent of cells expressing
c
Pou5f1
(Embryonic)
Plvap
(endocardium)
Tbx18
(epicardium)
Dlx5
(neural crest)
Max
Min

**Supplementary Fig. 5 | Detection of TEs in the embryonic heart scRNA-seq dataset.** (**a**) UMAP plot of embryonic heart scRNA-seq data, colored by embryonic stage (left) or by sample/replicate (right). (**b**) Expression of the indicated marker genes in the clusters as defined in Fig. 3i. Color indicates the normalized expression. The size of the dot indicates the percent of cells expressing that gene. (**c**) UMAP plots of selected marker genes, from the indicated lineages.

fig S6

a

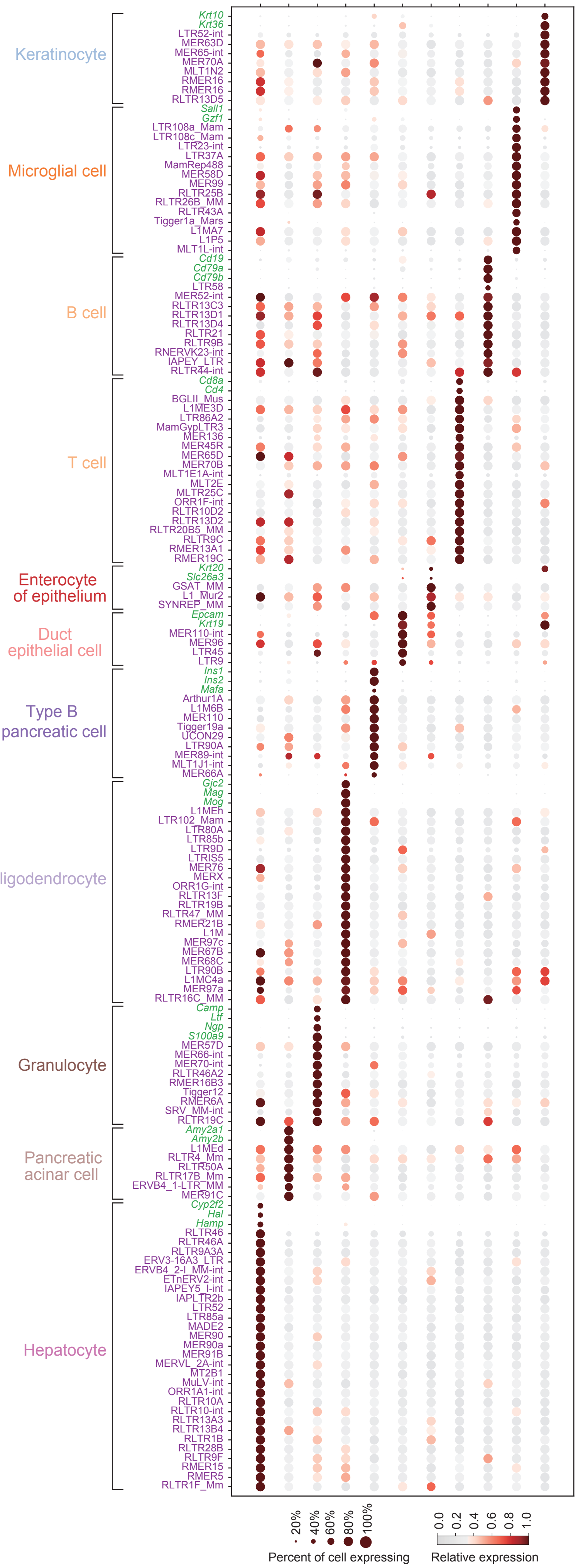


b

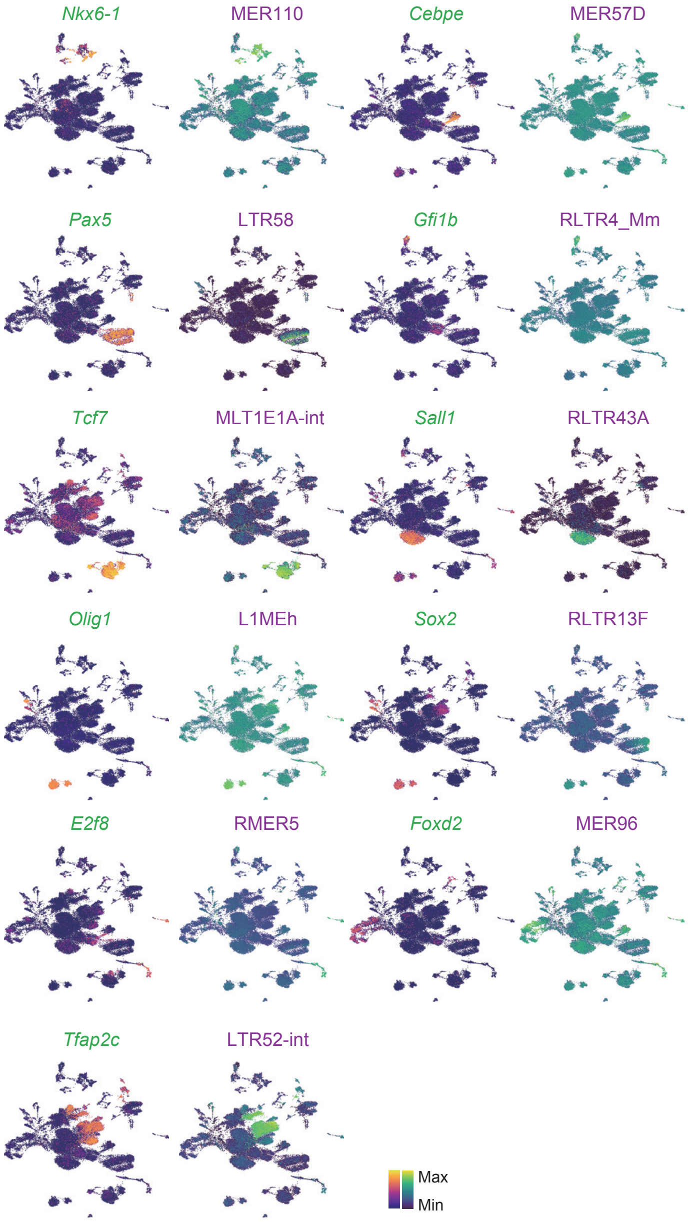


c

| p-value | TP Thresh | TP (%) | FP (%) |
|---|---|---|---|
| 7.16e-36 | 7.99 | 75.4 | 7.6 |

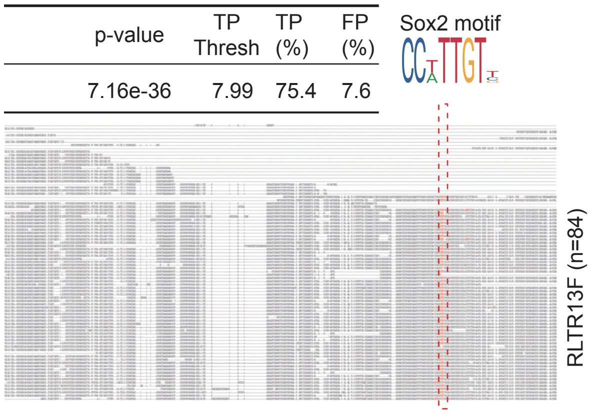

**Supplementary Fig. 6 I Cell-type specific expression of TEs.** (**a**) Dotplot showing the expression (color) and percentage of cells expressing the indicated marker genes for the indicated groups. (**b**) UMAP plots showing the expression of selected cell type-specific TFs andTEs. (**c**) Upper panel: Significantly enriched transcription factor binding motifs in the RLTR13F TEs. The motif analyses was measured using AME from the MEME suite; Lower panel: Alignment of all of the genomic copies of the RLTR13F, showing the location of the SOX2 motif in red. The consensus SOX2 sequence logo is indicated at the top of the TE. The number of copies of the TE are indicated (n=84).

fig S7

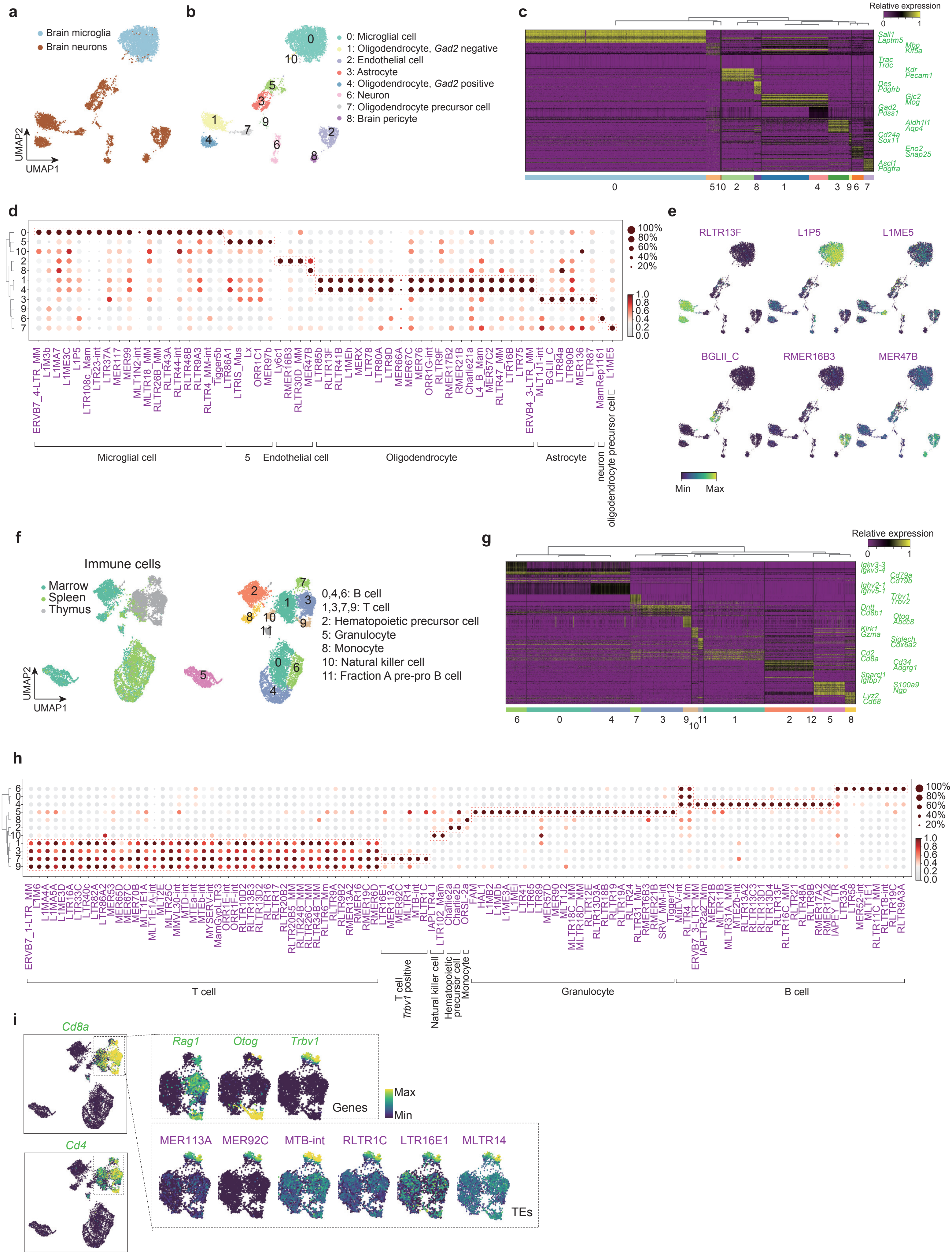
a
Brain microglia
Brain neurons
UMAP2
UMAP1
b
0: Microglial cell
1: Oligodendrocyte, Gad2 negative
2: Endothelial cell
3: Astrocyte
4: Oligodendrocyte, Gad2 positive
6: Neuron
7: Oligodendrocyte precursor cell
8: Brain pericyte
c
Relative expression
Sall1
Laptm5
Mbp
Kif5a
Trac
Trdc
Kdr
Pecam1
Des
Pdgfrb
Gjc2
Mog
Gad2
Pdss1
Aldh1l1
Aqp4
Cd24a
Sox11
Eno2
Snap25
Ascl1
Pdgfra
d
100%
80%
60%
40%
20%
Microglial cell
5
Endothelial cell
Oligodendrocyte
Astrocyte
neuron
oligodendrocyte precursor cell
e
RLTR13F
L1P5
L1ME5
BGLII_C
RMER16B3
MER47B
Min
Max
f
Immune cells
Marrow
Spleen
Thymus
0,4,6: B cell
1,3,7,9: T cell
2: Hematopoietic precursor cell
5: Granulocyte
8: Monocyte
10: Natural killer cell
11: Fraction A pre-pro B cell
UMAP2
UMAP1
g
Relative expression
Igkv3-3
Igkv3-4
Cd79a
Cd79b
Ighv2-1
Ighv5-1
Trbv1
Trbv2
Dntt
Cd8b1
Otog
Abcc8
Klrk1
Gzma
Siglech
Cox6a2
Cd2
Cd8a
Cd34
Adgrg1
Sparcl1
Igfbp7
S100a9
Ngp
Lyz2
Cd68
h
100%
80%
60%
40%
20%
T cell
T cell
Trbv1 positive
Natural killer cell
Hematopoietic precursor cell
Monocyte
Granulocyte
B cell
i
Cd8a
Cd4
Rag1
Otog
Trbv1
Genes
Max
Min
MER113A
MER92C
MTB-int
RLTR1C
LTR16E1
MLTR14
TEs

**Supplementary Fig. 7 I TEs are widely expressed in mouse brain and immune system**. (**a**) UMAP plot of mouse brain microglia and neurons. (**b**) UMAP plot as in panel a, but clustered into groups (Leiden, resolution=0.5). The indicated cell types are labelled according to the known marker genes from panel d. (**c**) A gene expression heatmap showing the top differentially expressed genes for each cell cluster from panel b. (**d**) Dotplot showing the expression (color) and percentage of cells expressing the indicated TEs, in the indicated groups from panel b. (**e**) UMAP plots showing the indicated TE expression across cell types. (**f**) UMAP plot of mouse marrow, spleen and thymus tissues. (**g**) A gene expression heatmap showing the top differentially expressed genes for each cell cluster as defined in panel f. Selected marker genes are indicated on the right-hand side. (**h**) Dot plot showing the expression (color) and percentage of cells expressing the indicated TEs, in the indicated groups from panel f. (**i**) UMAP plot showing the indicated gene and TE expression for T cell-specific genes/TEs

fig. S8

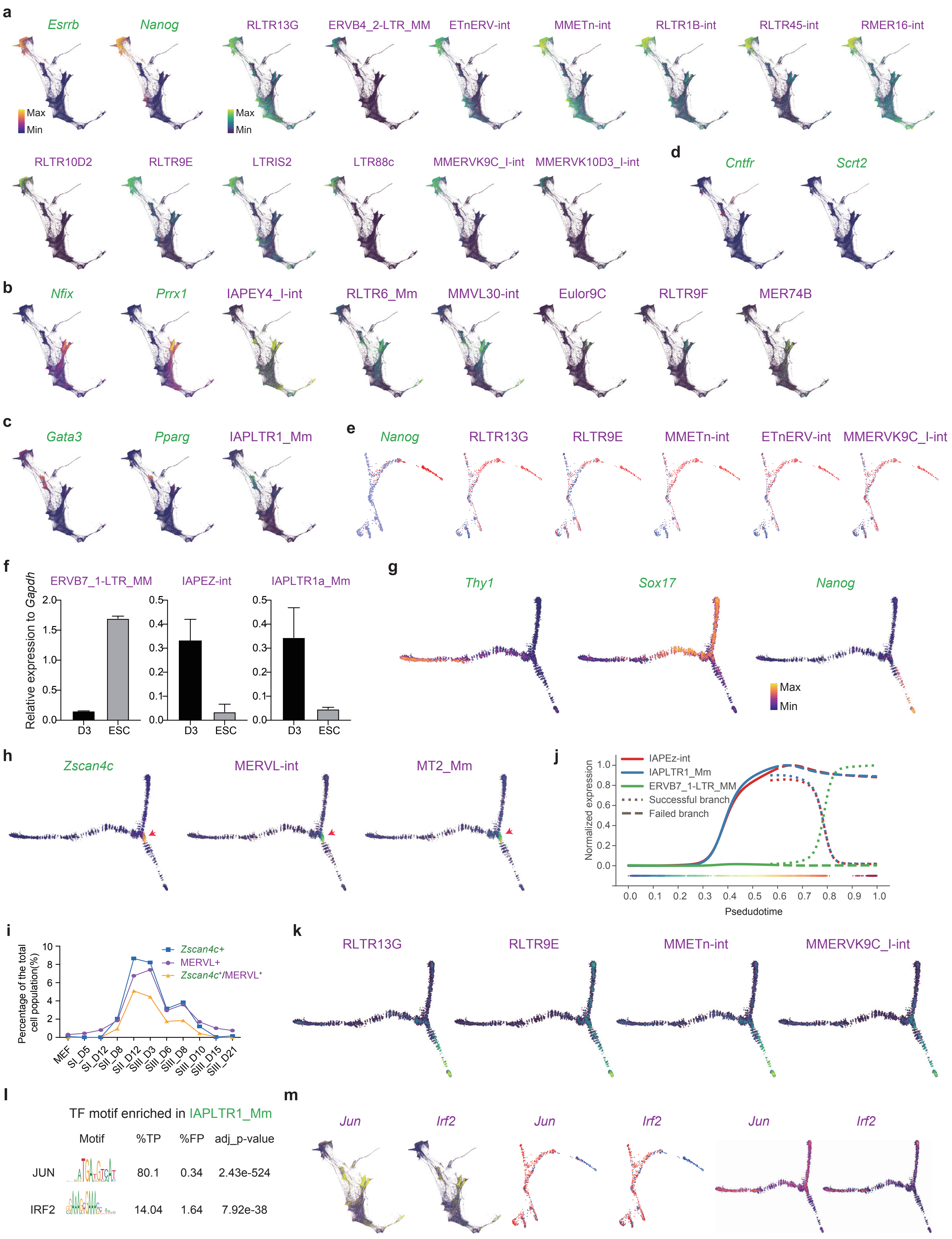
a
Esrrb
Nanog
RLTR13G
ERVB4_2-LTR_MM
ETnERV-int
MMETn-int
RLTR1B-int
RLTR45-int
RMER16-int
Max
Min
RLTR10D2
RLTR9E
LTRIS2
LTR88c
MMERVK9C_I-int
MMERVK10D3_I-int
d
Cntfr
Scrt2
b
Nfix
Prrx1
IAPEY4_I-int
RLTR6_Mm
MMVL30-int
Eulor9C
RLTR9F
MER74B
c
Gata3
Pparg
IAPLTR1_Mm
e
Nanog
RLTR13G
RLTR9E
MMETn-int
ETnERV-int
MMERVK9C_I-int
f
Relative expression to Gapdh
ERVB7_1-LTR_MM
IAPEZ-int
IAPLTR1a_Mm
D3
ESC
g
Thy1
Sox17
Nanog
Max
Min
h
Zscan4c
MERVL-int
MT2_Mm
j
IAPEz-int
IAPLTR1_Mm
ERVB7_1-LTR_MM
Successful branch
Failed branch
Normalized expression
Psedudotime
i
Percentage of the total cell population(%)
Zscan4c+
MERVL+
Zscan4c+/MERVL+
MEF
SI_D5
SI_D12
SII_D8
SII_D12
SIII_D3
SIII_D6
SIII_D8
SIII_D10
SIII_D15
SIII_D21
k
RLTR13G
RLTR9E
MMETn-int
MMERVK9C_I-int
l
TF motif enriched in IAPLTR1_Mm
Motif
%TP
%FP
adj_p-value
JUN
80.1
0.34
2.43e-524
IRF2
14.04
1.64
7.92e-38
m
Jun
Irf2
Jun
Irf2
Jun
Irf2

**Supplementary Fig. 8 | Dynamic expression of TEs during somatic cell reprogramming.** (**a**) Visualization of scRNA-seq profiles for OKSM-based reprogramming, cells are colored by the expression of pluripotency marker genes and selected TEs. (**b**) Expression of stromal marker genes and selected TEs. (**c**) Expression of trophoblast marker genes and the TE IAPLTR1_Mm. (**d**) Expression of neural marker genes. (**e**) Expression of the pluripotent marker gene *Nanog* or TEs during OKS reprogramming. (**f**) Bar plot show the qRT-PCR analysis the expression of selected TEs. Primers used are described in table S1. (**g**) Expression of marker genes during chemical reprogramming. (**h**) As in panel g, but showing the expression of 2C-like related genes or TEs. (**i**) Percent of 2C-like cells at different time points during chemical reprogramming. (**j**) Expression kinetics of selected maker TEs along pseudotime during reprogramming. (**k**) Expression dynamics of the indicated TEs during chemical reprogramming. (**l**) Significantly enriched transcription factor binding motifs in the IAPLTR1_Mm TEs. The motif analyses were measured using AME from the MEME suite. (**m**) Expression level of *Jun* and *Irf2* in the three different reprogramming systems, OKSM (left), OKS (middle) and chemical-reprogramming (right).

### fig. S9

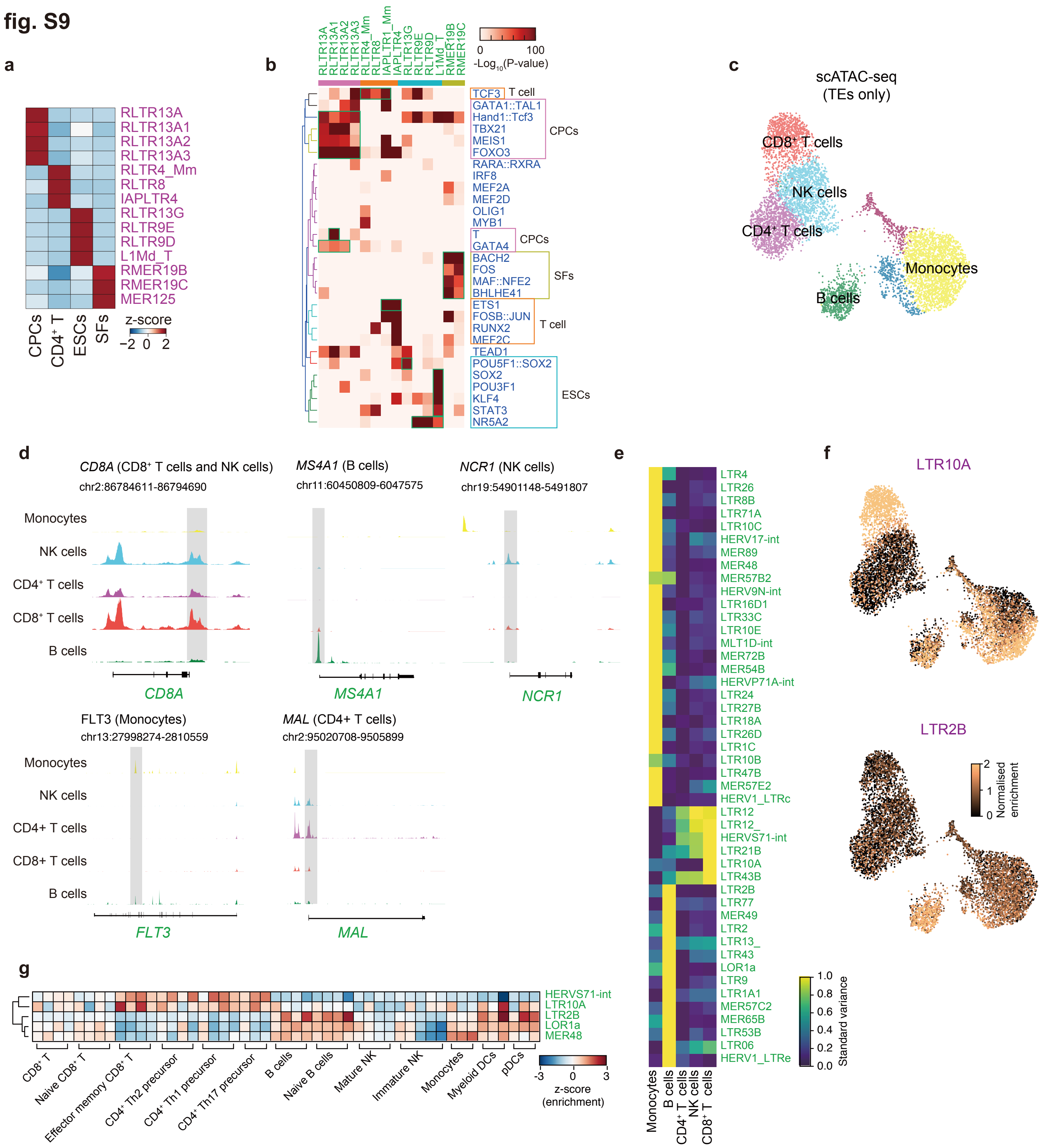

a
RLTR13A
RLTR13A1
RLTR13A2
RLTR13A3
RLTR4_Mm
RLTR8
IAPLTR4
RLTR13G
RLTR9E
RLTR9D
L1Md_T
RMER19B
RMER19C
MER125
CPCs
CD4+ T
ESCs
SFs
z-score
−2 0 2
b
0 100
-Log10(P-value)
TCF3 T cell
GATA1::TAL1
Hand1::Tcf3
TBX21
MEIS1
FOXO3
CPCs
RARA::RXRA
IRF8
MEF2A
MEF2D
OLIG1
MYB1
T
GATA4
CPCs
BACH2
FOS
MAF::NFE2
BHLHE41
SFs
ETS1
FOSB::JUN
RUNX2
MEF2C
T cell
TEAD1
POU5F1::SOX2
SOX2
POU3F1
KLF4
STAT3
NR5A2
ESCs
c
scATAC-seq
(TEs only)
CD8+ T cells
NK cells
CD4+ T cells
Monocytes
B cells
d
CD8A (CD8+ T cells and NK cells)
chr2:86784611-86794690
MS4A1 (B cells)
chr11:60450809-6047575
NCR1 (NK cells)
chr19:54901148-5491807
Monocytes
NK cells
CD4+ T cells
CD8+ T cells
B cells
CD8A
MS4A1
NCR1
FLT3 (Monocytes)
chr13:27998274-2810559
MAL (CD4+ T cells)
chr2:95020708-9505899
FLT3
MAL
e
LTR4
LTR26
LTR8B
LTR71A
LTR10C
HERV17-int
MER89
MER48
MER57B2
HERV9N-int
LTR16D1
LTR33C
LTR10E
MLT1D-int
MER72B
MER54B
HERVP71A-int
LTR24
LTR27B
LTR18A
LTR26D
LTR1C
LTR10B
LTR47B
MER57E2
HERV1_LTRc
LTR12
LTR12_
HERVS71-int
LTR21B
LTR10A
LTR43B
LTR2B
LTR77
MER49
LTR2
LTR13_
LTR43
LOR1a
LTR9
LTR1A1
MER57C2
MER65B
LTR53B
LTR06
HERV1_LTRe
Standard variance
1.0 0.8 0.6 0.4 0.2 0.0
Monocytes
B cells
CD4+ T cells
NK cells
CD8+ T cells
f
LTR10A
LTR2B
Normalised enrichment
2 1 0
g
HERVS71-int
LTR10A
LTR2B
LOR1a
MER48
CD8+ T
Naive CD8+ T
Effector memory CD8+ T
CD4+ Th2 precursor
CD4+ Th1 precursor
CD4+ Th17 precursor
B cells
Naive B cells
Mature NK
Immature NK
Monocytes
Myeloid DCs
pDCs
-3 0 3
z-score
(enrichment)

**Supplementary Fig. 9 | Analysis of the Single-Cell ATAC-seq data**. (**a**) Heatmap showing the enrichment of selected TEs across cell types in pseudo-bulk ATAC-seq data generated from the scRNA-seq. (**b**) Heatmap showing TF motif enrichment in selected TEs. (**c**) UMAP plot of the TE chromatin state from PBMC scATAC-seq data, cells were colored by cell types. (Leiden clustering, resolution=0.8). Cell types were annotated based on the specific opened marker genes (See panel D). Data was from 10x genomics website. (**d**) Genome track plots showing the aggregated scATAC-seq profiles of selected marker genes for the indicated lineages. (**e**) Heatmap of significantly differentially open TEs between the indicated cell types. (Benjamini-Hochberg corrected Wilcoxon rank-sum test, p-value<0.01), and at least >2 fold change between groups. (**f**) UMAP plot, as panel C, but cells are colored by expression of the indicated TEs. (**g**) Heatmap showing a z-score enrichment of selected TEs across different immune cell types from bulk ATAC-seq data.

fig.S10

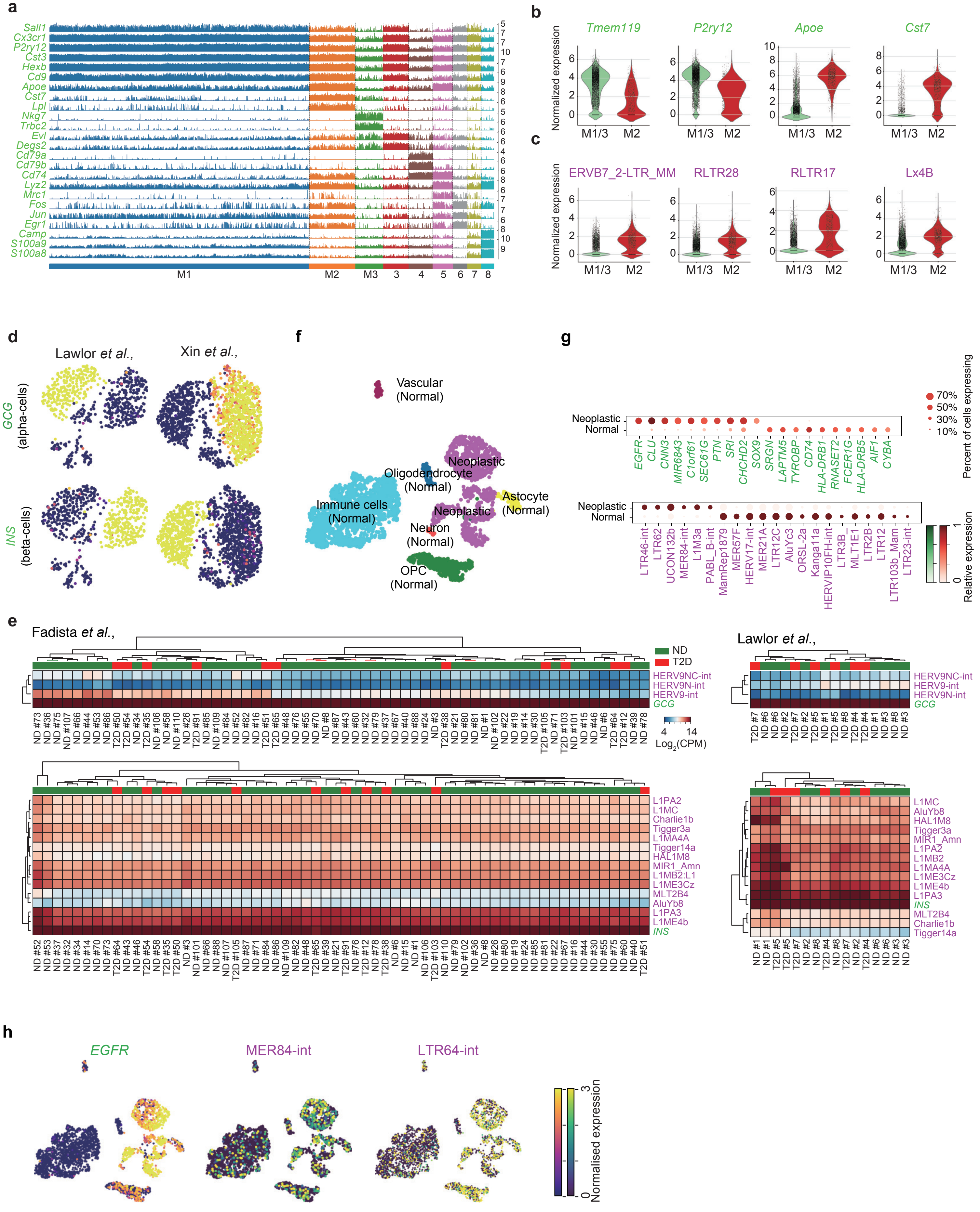
a
Sall1
Cx3cr1
P2ry12
Cst3
Hexb
Cd9
Apoe
Cst7
Lpl
Nkg7
Trbc2
Evl
Degs2
Cd79a
Cd79b
Cd74
Lyz2
Mrc1
Fos
Jun
Egr1
Camp
S100a9
S100a8
M1
M2
M3
b
Tmem119
P2ry12
Apoe
Cst7
Normalized expression
M1/3
M2
c
ERVB7_2-LTR_MM
RLTR28
RLTR17
Lx4B
d
Lawlor et al.,
Xin et al.,
GCG
(alpha-cells)
INS
(beta-cells)
f
Vascular
(Normal)
Neoplastic
Oligodendrocyte
(Normal)
Immune cells
(Normal)
Astocyte
(Normal)
Neuron
(Normal)
OPC
(Normal)
g
Neoplastic
Normal
EGFR
CLU
CNN3
MIR6843
C1orf61
SEC61G
PTN
SRI
CHCHD2
SOX9
SRGN
LAPTM5
TYROBP
CD74
HLA-DRB1
RNASET2
FCER1G
HLA-DRB5
AIF1
CYBA
70%
50%
30%
10%
Percent of cells expressing
LTR46-int
LTR62
UCON132b
MER84-int
L1M3a
PABL_B-int
MamRep1879
MER57F
HERV17-int
MER21A
LTR12C
AluYc3
ORSL-2a
Kanga11a
HERVIP10FH-int
LTR3B_
MLT1E1
LTR2B
LTR12
LTR103b_Mam
LTR23-int
Relative expression
e
Fadista et al.,
ND
T2D
HERV9NC-int
HERV9N-int
HERV9-int
GCG
Log2(CPM)
Lawlor et al.,
HERV9NC-int
HERV9-int
HERV9N-int
L1PA2
L1MC
Charlie1b
Tigger3a
L1MA4A
Tigger14a
HAL1M8
MIR1_Amn
L1MB2:L1
L1ME3Cz
MLT2B4
AluYb8
L1PA3
L1ME4b
INS
L1MC
AluYb8
HAL1M8
Tigger3a
MIR1_Amn
L1PA2
L1MB2
L1MA4A
L1ME3Cz
L1ME4b
L1PA3
INS
MLT2B4
Charlie1b
Tigger14a
h
EGFR
MER84-int
LTR64-int
Normalised expression

**Supplementary Fig. 10 | TE and marker gene expression in a mouse model of Alzheimer's disease, in human type 2 diabetes and glioblastoma.** (**a**) Tracksplot showing the known marker gene expression for cell clusters, the cell clusters are from Fig. 6b. The x-axis is for cells, each column represents an individual cell, and the y-axis represent the expression value. (**b**) Violin plots showing the expression of known marker genes. (**c**) Violin plots, as in panel b, showing the expression of indicated TEs. (**d**) UMAP plots showing the indicated marker gene expression for alpha and beta cells. The yellow represents high expressed and black represent low expressed. (**e**) Heatmap of expression for the indicated genes and TEs in bulk RNA-seq data. The '#' number indicates the patient, and T2D=type 2 diabetes and ND=Non-diabetic. (**f**) UMAP plot of human glioblastoma, the metadata provided in the original study was used to label the cell identity. (**g**) Dot plot showing the differentially expressed genes and TEs between normal and neoplastic group cells. (**h**) UMAP plots showing the expression of selected TEs and *EGFR*, a known glioblastoma neoplastic gene.
